# Identification and characterisation of *Planococcus citri cis*- and *trans*-isoprenyl diphosphate synthase genes, supported by short- and long-read transcriptome data

**DOI:** 10.1101/2023.06.09.544309

**Authors:** Mojca Juteršek, Iryna M. Gerasymenko, Marko Petek, Elisabeth Haumann, Sandra Vacas, Kalyani Kallam, Silvia Gianoglio, Vicente Navarro-Llopis, Ismael Navarro Fuertes, Nicola Patron, Diego Orzáez, Kristina Gruden, Heribert Warzecha, Špela Baebler

**Author notes:** These authors contributed equally to the work (Mojca Juteršek, Iryna M. Gerasymenko). Coresspondence to.

## Abstract

Many insect species rely on diverse terpenoids for their development and interorganismal interactions. However, little is known about terpenoid biosynthesis in insects. The monoterpenoid sex pheromones of mealybugs and scale insects (Coccoidea) are particularly enigmatic, with several species producing unique structures presumed to result from the irregular coupling activity of unidentified isopentenyl diphosphate synthases (IDSs). Enzymes capable of similar transformations have previously only been described from a few plant, bacterial and archaeal species. To investigate if insect irregular monoterpenes can be biosynthesised by similar enzymes, we performed a comprehensive search for IDS coding sequences in the genome of *Planococcus citri*, a widespread agricultural pest. We complemented the available *P. citri* genome data with newly generated short- and long-read transcriptome data. The identified candidate genes had homology to both short- and long-chain IDSs and some appeared to be paralogous, indicating gene duplications and consequent IDS gene family expansion in *P. citri*. We tested the activity of eleven candidate gene products, confirming *in vitro* regular activity for five enzymes, one of which (*trans*IDS5) also produced the irregular prenyl diphosphates, maconelliyl and lavandulyl diphosphate. Targeted mutagenesis of selected aspartates and a lysine in the active site of *trans*IDS5 uncovered their importance for chain-length preference and irregular coupling. This work provides an important foundation for deciphering terpenoid biosynthesis in mealybugs, as well as a potential source of enzymes for the biotechnological production of sustainable insect pest management products.

## Introduction

Terpenoids are a large class of metabolites produced by organisms in every branch of life. In insects, terpenoids have important roles in development (e.g., juvenile and moulting hormones) and as semiochemicals in intra-as well as interspecific interactions (e.g., sex, aggregating, alarm, dispersal, maturation, anti-aphrodisiac or trail pheromones, and defence compounds) (Tillman et al., 1999; Beran et al., 2019). Of special interest are the terpenoid sex pheromones produced by some mealybug and scale insect species (order Hemiptera, Superfamily Coccoidea). Sex pheromones are species-specific, volatile, long-range attractants. In Coccoidea, they are produced by females to facilitate male navigation during mating and are thus emitted by virgin females, with cessation of production after mating (Zou and Millar, 2015). The majority of mealybug (Family Pseudococcidae) and armoured scale insect (Family Diaspididae) sex pheromones identified to date are irregular terpenoid compounds, mostly esterified mono- or sesquiterpenoid alcohols and carboxylic acids (Zou and Millar, 2015; Franco et al., 2022). Chemically synthesised mealybug and scale insect pheromones have been successfully used for pest control, providing a sustainable measure in climate change-challenged agriculture (Vacas et al., 2010; Zou et al., 2013; Zou and Millar, 2015; Lucchi et al., 2019; Daane et al., 2021). Although an extensive body of research has been dedicated to the identification and study of mealybug and scale insect sex pheromones, the biosynthesis of their irregular backbone remains elusive.

The first step of terpenoid biosynthesis is catalysed by isoprenyl diphosphate synthases (IDSs or prenyltransferases), which couple the C5 terpene building blocks, isopentenyl diphosphate (IPP) and dimethylallyl diphosphate (DMAPP) into C10 (geranyl diphosphate, GPP), C15 (farnesyl diphosphate, FPP), C20 (geranylgeranyl diphosphate, GGPP) and longer prenyl diphosphate precursor chains by sequential condensation steps (Nagel et al., 2019). The prenyl diphosphate chains are then transformed into terpenoid compounds through the activity of terpene synthases (TPSs).

IDSs can be classified as either *cis*- or *trans*-, depending on the stereochemistry of the double bond in the reaction product. The two classes are evolutionarily and structurally distinct. *Trans*-IDSs adopt an α-fold and contain two conserved aspartate-rich motifs known as FARM (first aspartate rich motif) and SARM (second aspartate rich motif), which mainly occur as “DDxx(x)D”. *Cis*-IDSs, however, lack distinct conserved motifs and adopt the ζ-fold (Nagel et al., 2019). Aspartate-rich motifs in *trans*-IDSs coordinate the Mg^2+^ ions important for generation of the carbocation in the allylic substrate (DMAPP, GPP, FPP or other), which can be attacked by IPP to form the new carbon-carbon (C-C) bond in the alkylation reaction. The coupling step catalysed by IDSs can be also classified based on the orientation of the alkylation reaction. Most IDS enzymes catalyse the formation of regular 1’-4 (head-to-tail) C-C bond between the allylic substrate and IPP, while irregular head-to-middle reactions have been described for some *cis*- and *trans*-IDS enzymes, coupling two DMAPP units or DMAPP with an allylic substrate, forming branched or cyclic polyprenyl chains (Kobayashi and Kuzuyama, 2018). To date, only a few enzymes with irregular coupling activity have been identified from plants (Rivera et al., 2001; Demissie et al., 2013), bacteria (Ozaki et al., 2014), and archaea (Ogawa et al., 2016).

*Planococcus citri* (Risso, 1813), the citrus mealybug, is a small sucking insect. It is a widespread pest of numerous crops and ornamental plants, causing serious damage and economic losses (Watson, 2023). Its sex pheromone, planococcyl acetate, has an cyclobutane backbone, presumably resulting from irregular IDS coupling. It has been successfully chemically synthesised and deployed in integrated pest management via trap-based management strategies, which require only milligram quantities (Bierl-Leonhardt et al., 1981; Wolk et al., 1986; Dunkelblum et al., 2002; Passaro and Webster, 2004). Alternative management strategies, such as mating disruption, demand larger quantities and, therefore, require low-cost, scalable production methods (Zou and Millar, 2015). Biotechnological production of sex pheromones has been demonstrated recently for lepidopteran pheromones (Mateos-Fernández et al., 2021; Petkevicius et al., 2022) and biotechnological synthesis has been proposed as a sustainable route for large-scale, low-cost pheromone production. However, these methods require knowledge of the biosynthetic pathways.

To that end, we performed a comprehensive search of the *P. citri* genome, supported by newly generated short- and long-read sequence datasets of *P. citri* females, to identify candidate *cis*- and *trans*-IDSs. Eleven of the identified IDS sequences were tested for regular and irregular coupling activity, with a focus on the production of short-chain backbones. We confirmed regular coupling activity for five *trans*-IDS candidate enzymes, one of which also produced smaller amounts of the irregular lavandulyl and maconelliyl diphosphates. Our work contributes to understanding of mealybug terpenoid biosynthesis providing a foundation for its biotechnological exploitation.

## Results

### *Planococcus citri* short- and long-read transcriptome resource

To complement the available *P. citri* genomic sequence data (*Pcitri*.v1 genome assembly) we generated short and long transcriptome reads by performing RNA-Seq and Iso-Seq. Short Illumina reads from virgin and mated *P. citri* females generated in this study were *de novo* assembled on its own (Figure 1a), and together with other available *P. citri* short-read sequence data, including reads from *P. citri* males (Figure 1b). Virgin and mated female-specific transcriptomic data was generated to obtain information on possible pheromone synthesis-related changes in gene expression. Both assemblies were combined, resulting in 389,962 unique sequences (Figure 1c). After the merge with the long-read transcriptome (Figure 1d), the full consolidated set contained 440,881 unique transcript sequences (Figure 1e), with an average length of 1,356 nucleotides and N50 of 2,908 nucleotides (Table S1). On average, around 80 % of short reads from each sample mapped back to *de novo* transcriptome assembly (Table S2). To retain as much variability as possible, we did not do any further assembly thinning and the presented dataset is, therefore, redundant. Approximately 53 % of the transcripts mapped to the *Pcitri*.v1 genome, with a third of them mapping to the annotated *Pcitri*.v1 gene models. Completeness assessment with BUSCO found 98.7 % complete BUSCOs for the consolidated transcriptome, the majority of which were duplicated (Figure S1). Transcript IDs with their InterPro annotations, information on mapping to *Pcitri*.v1 genome and virgin versus mated female differential expression values are available in Table S3a, as well as on FAIRDOMHub along with a fasta file with all sequences from the consolidated transcriptome (see Data Availability Statement).

**Figure 1:**
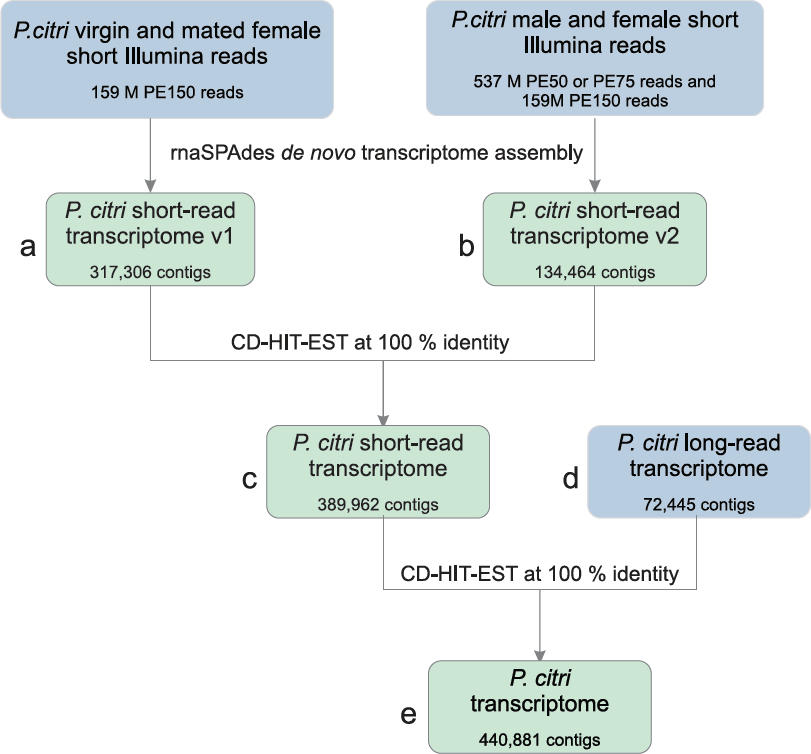
Construction of *P. citri* transcriptome resource. Diagram of consolidation steps taken to combine the short-read (a, b, merged into c) and long-read (d) *P. citri* transcriptome resources into a comprehensive dataset of all assembled and sequenced putative *P. citri* transcripts (e). Raw sequencing data are in blue boxes.

Due to the elusive nature of terpenoid biosynthesis in insects, it has been hypothesised that the origin of some terpenoids (or their precursors), might not be the insects themselves, but rather endosymbiotic bacteria or even plant food (Beran and Petschenka, 2022). For a comprehensive analysis of the biosynthetic capacity for terpenoids found in *P. citri*, our transcriptomic resource, therefore, contains transcripts of all taxonomic origins, as we elected not to exclude non-insect reads. Taxonomic analyses revealed that out of 207,753 protein sequences determined from the consolidated transcriptome, 26 % were unclassified, with the majority of classified sequences assigned to insects (Figure S2). 2,641 sequences were assigned to bacteria, 1,010 of which correlated best with sequences from *Candidatus* Moranella endobia, a secondary endosymbiont of *P. citri*. The acquired data also includes 2,364 viral, 1,629 plant, and 4,299 fungal sequences. Of the plant sequences, most were assigned to potato (*Solanum tuberosum*), probably originating from the *P. citri* food source. Of the fungal sequences, most were assigned to *Wallemia mellicola*, a cosmopolitan fungus with air-disseminated spores (Jančič et al., 2015).

### Identification of putative IDS coding sequences

To find sequences with homology to *cis*- and *trans*-IDS enzymes, we searched *Pcitri*.v1 genome assembly as well as the *P. citri* consolidated transcriptome (Figure 1e) for sequences with assigned InterPro family annotations IPR001441, and IPR036424 (for *cis*-IDS sequences) or IPR000092 (for *trans*-IDS sequences). On the *Pcitri*.v1 genome assembly we also performed MAST motif search. All extracted sequences (Table S4a and b) were collapsed into clusters based on their sequence similarity. This resulted in a final number of 30 putative IDS sequences (Table S4c). Newly acquired transcript sequence information provided alternative primary sequences to matching *Pcitri*.v1 gene models for seven IDS sequences (Figure 2, Figure S3). For five of them (*cis*IDS1, *trans*IDS4, *trans*IDS5, *trans*IDS11, and *trans*IDS12) we were able to confirm the sequences predicted by the transcriptome data with amplification (Figure 2).

**Figure 2:**
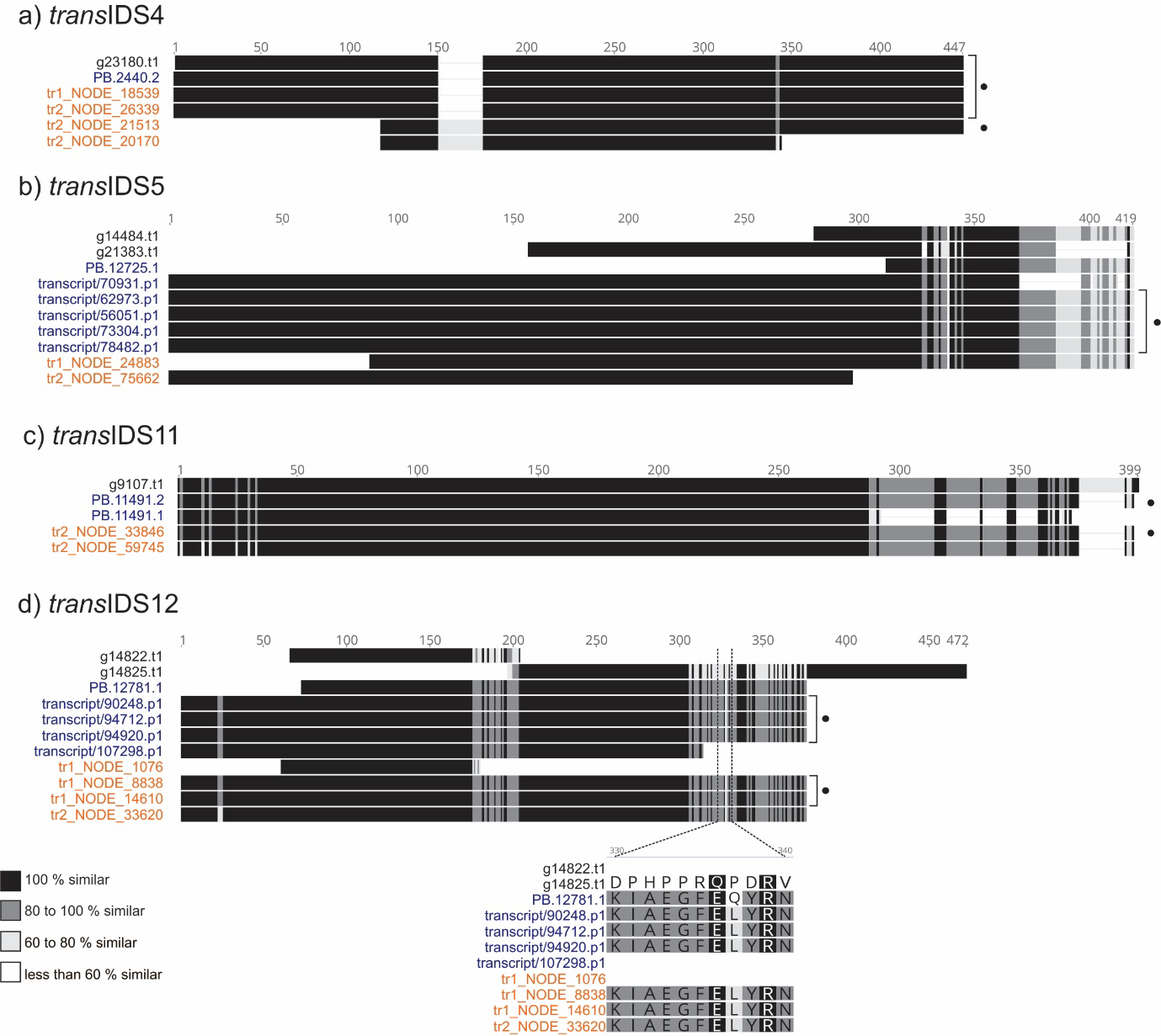
Identification and confirmation of IDS coding sequences predicted by transcriptome data. Multiple sequence alignments for clusters of overlapping transcripts and gene models for candidate sequences *trans*IDS4 (a), *trans*IDS5 (b), *trans*IDS11 (c), and *trans*IDS12 (d). Each cluster includes translated coding sequences obtained from *Pcitri*.v1 gene models (in black text), long-read transcripts (in blue text), and transcripts *de novo* assembled from short reads (in orange text). All sequences are available in Table S4. Sequences confirmed with amplification from cDNA and Sanger sequencing are marked with black dots. Colour-coded sequence similarity is given in the legend in the bottom left. The translated amplicon of *cis*IDS1 has Leu (L327) instead of His at position 327 (H327), the latter predicted in the gene model, while the transcripts all have L327 (not shown in Figure). Two different amplicons were obtained in the case of *trans*IDS4 (a), one identical to the gene model and one with N-terminal truncation and an insertion of 25 amino acids. For *trans*IDS5 (b), the gene model was missing 283 amino acids at the N-terminus. For *trans*IDS11 (c), we amplified a sequence with a frame-shift close to the C-terminus resulting in a 21 amino acids long C-terminus truncation. In the case of *trans*IDS12 (d), two variations of the same full-length sequence were amplified, differing in a single amino acid position: L328 versus Q328. The alignment around the L328 versus Q328 variation is shown below the full length alignments. The structure of the amplified sequence of *trans*IDS12 differed significantly from the annotated gene models. Multiple sequence alignment was done in MEGAX using the MUSCLE algorithm and visualised with Geneious software.

Eight of the identified IDS sequences most likely originated from other species: *trans*IDS14 has only one mismatch to a FPPS from potato, while *trans*IDS15, *cis*IDS7, and *cis*IDS10 have 100 % identity to different IDS sequences from the fungus *Wallemia mellicola*, respectively (Table S4c). *Trans*IDS6, *trans*IDS7, *cis*IDS4, and *cis*IDS6 have homology to insect IDS sequences. However, as determined by phylogenetic analysis (Supplementary Figures S4, S5, S6, and S7), these grouped with homologous sequences from Chalcidoidea wasps (Hymenoptera). Additionally, one sequence can be assigned to a *P. citri* symbiont: *cis*IDS8 is identical to a protein annotated as a UPPS (undecaprenyl diphosphate synthase) from *Candidatus* Moranella endobia.

### *In silico* analysis reveals diverse putative functions of candidate *P. citri* IDSs

We further analysed the sequence features and phylogeny of identified *P. citri* IDS sequences, which suggested their potential functions (Table 1). Two candidate sequences (*trans*IDS2, *trans*IDS4) are similar to decaprenyl diphosphate synthases (DPPSs), long-chain IDSs that catalyse the formation of the ubiquinone prenyl sidechain, important in aerobic cellular respiration. The functional DPPS enzyme forms a heterodimer of subunits 1 and 2, of which subunit 1 has a higher sequence homology to IDSs, while subunit 2 acts as a regulatory subunit (Zhang and Li, 2013; Song and Li, 2022). T*rans*IDS2 is similar to sequences of DPPS subunit 1, while *trans*IDS4 lacks the D-rich motifs and is similar to sequences of DPPS subunit 2 (Table 1, Supplementary Figure S4, Supplementary Figure S8).

**Table 1:** *P. citri cis*- and *trans*-IDS candidates. For each candidate sequence, its putative function, differential expression values (logFC), contrasting samples from virgin to mated females, and results of *in vitro* regular (R) and irregular (I) C10 or C15 coupling activity tests of the proteins expressed in *E. coli* are given. For *trans*-IDS candidates, amino acids positioned at the first and second D-rich motif (FARM, SARM) according to the multiple sequence alignment are given as well. LogFC (log2-fold changes) values were obtained by averaging logFC values for all sequences in the sequence cluster of each candidate (Table S4c) and are coloured in red and green (for up- and downregulated expression, respectively) if the differential expression was statistically significant (pAdj > 0.05) for at least one of the sequences in each cluster. “/” – activity test not performed. Asterisk denotes the sequence originating from *Candidatus* Moranella endobia.

| Candidate | FARM | SARM | Putative function | logFC | R | I |
| --- | --- | --- | --- | --- | --- | --- |
| *trans*IDS2 | DDVID | DDLLD | DPPS subunit 1 | 0.18 | Y | N |
| *trans*IDS3 | DDIID | DDYLD | FPPS | 1.20 | Y | N |
| *trans*IDS4 | / | / | DPPS subunit 2 | 0.32 | N | N |
| *trans*IDS5 | DDIMD | DDYLD | FPPS | 0.22 | Y | Y |
| *trans*IDS8 | DDILD | NDYND | FPPS-like | -1.26 | / | / |
| *trans*IDS9 | DDIFD | NDYND | FPPS-like | 3.11 | / | / |
| *trans*IDS10 | DDAID | KDQND | FPPS-like | 0.93 | N | N |
| *trans*IDS11 | DDALD | NDYYD | FPPS-like | 1.61 | Y | N |
| *trans*IDS12 | DDIAD | KLYND | FPPS-like | 5.49 | N | N |
| *trans*IDS13 | DDVVD | KIFND | FPPS-like | -1.22 | / | / |
| *trans*IDS16 | DDIQD | DDYCN | GGPPS | 0.08 | N | N |
| *trans*IDS17 | DDFHD | / | ubiquitin thioesterase | -0.13 | Y | N |
| *trans*IDS18 | DDALD | NDYYD | FPPS-like | -0.38 | / | / |
| *cis*IDS1 | / | / | DHPPS | 0.07 | N | N |
| *cis*IDS2 | / | / | DHPPS-like | 1.90 | / | / |
| *cis*IDS3 | / | / | DHPPS-like | 0.09 | / | / |
| *cis*IDS5 | / | / | DHPPS-like | 0.04 | / | / |
| *cis*IDS8* | / | / | UPPS | NA | N | N |
| *cis*IDS9 | / | / | DHPPS regulatory subunit | 0.15 | / | / |

*Trans*IDS3 and *trans*IDS5 have high similarity to FPPSs (Table 1, Supplementary Figure S5). Additionally, there are seven sequences (*trans*IDS8, *trans*IDS9, *trans*IDS10, *trans*IDS11, *trans*IDS12, *trans*IDS13, *trans*IDS18) with similarity to FPPS-like sequences from other Coccoidea species (Supplementary Figure S9). All seven of them have a canonical FARM (DDxxD), but not SARM (Table 1). Phylogenetically, the FPPS-like sequences are separated from the mealybug FPPS sequences with low sequence similarity between the two groups (Supplementary Figure S9). Similar diversification pattern of IDS genes within Coccomorpha has been indicated previously (Rebholz et al., 2023). *Trans*IDS3 and *trans*IDS5, as well as *trans*IDS12 and *trans*IDS13 also have high pairwise sequence similarity (78 % and 75 %, respectively), indicating a possibility of more recent duplication events.

*Trans*IDS16 is similar to geranylgeranyl pyrophosphate synthases (GGPPS), catalysing the formation of the diterpenoid precursor (Supplementary Figure S10). Sequences of *trans*IDS1, *trans*IDS19, and *trans*IDS20 do not have canonical D-rich motifs. Additionally, *trans*IDS1 is similar to E3 ubiquitin protein ligases, while we could not find any similarity matches for *trans*IDS19 and *trans*IDS20. *Trans*IDS17 is similar to otubain-like ubiquitin thioesterases (Supplementary Figure S11) and includes a DDxxD motif aligning with FARM, as do all other otubain-like orthologs identified from mealybugs and included in the phylogenetic analysis (Supplementary Figure S11).

Out of the *cis*-IDS candidates, *cis*IDS1, *cis*IDS2, *cis*IDS3, and *cis*IDS5 are homologous to the catalytic subunit of heterodimeric dehydrodolichyl diphosphate synthase (DHPPS) (Supplementary Figure S6, Supplementary Figure S12), involved in the biosynthesis of dolichol phosphate, the lipid carrier of glycosyl units used for N-glycosylation in eukaryotes. Candidates *cis*IDS2, *cis*IDS3, and *cis*IDS5 have some closer orthologs within the Coccoidea, however they are very distant to other DHPPS sequences and even *cis*IDS1 (Supplementary Figure S12). On the other hand, *cis*IDS9 has highest similarity to sequences of the DHPPS regulatory subunit (Supplementary Figure S7).

### Regular IDS activity was shown for five and irregular activity for one *P. citri* candidate

With the aim of identifying enzymes involved in the synthesis of the irregular C10 sex pheromone, we tested the activity of *P. citri* IDS candidates for producing short polyprenyl chains. We selected 11 *P. citri* candidate sequences with homology to *trans*- or *cis*-IDSs for which the full sequence was confirmed with long-read transcriptome sequencing (Table 1, Table S4c). Two of these sequences (*trans*IDS3 and *trans*IDS10) were synthesised according to long-read sequencing data, and nine sequences (*trans*IDS2, *trans*IDS4, *trans*IDS5, *trans*IDS11, *trans*IDS12, *trans*IDS16, *trans*IDS17, *cis*IDS1, and *cis*IDS8) were amplified from *P. citri* cDNA (Table S4c). All sequences were expressed in *E. coli* and purified recombinant proteins were tested *in vitro* to identify regular and irregular IDS catalytic activity forming C10 or C15 backbones.

Five *trans*-IDS-like proteins (*trans*IDS2, *trans*IDS3, *trans*IDS5, *trans*IDS11, *trans*IDS17) displayed regular activity (Table 1, Figure 3a). The products of *trans*IDS2, *trans*IDS5, *trans*IDS11, and *trans*IDS17 were identified as geranyl and farnesyl diphosphates based on chromatographic behaviour and mass spectra of diphosphates and their dephosphorylated alcohol derivatives (Figure 3b, Figures S13-S19). Candidate *trans*IDS3 had an unusual pattern of regular products. It formed two isomers of C10 prenyl diphosphate, geranyl and *iso*-geranyl diphosphates, that were further elongated to two C15 isomers: farnesyl and *iso*-farnesyl diphosphate (Figure 3b). Specific activities of *trans*IDS5 were significantly higher than those of the other enzymes, with *trans*IDS3 also exhibiting a relatively high activity for production of C10 prenyl diphosphates (Figure 3a, Table S5). Both *trans*IDS3 and *trans*IDS5 sequences were expressed in *P. citri* females with *trans*IDS3 also differentially expressed between virgin and mated females with higher expression in virgin females (Table 1, Supplementary Table S4c).

**Figure 3:**
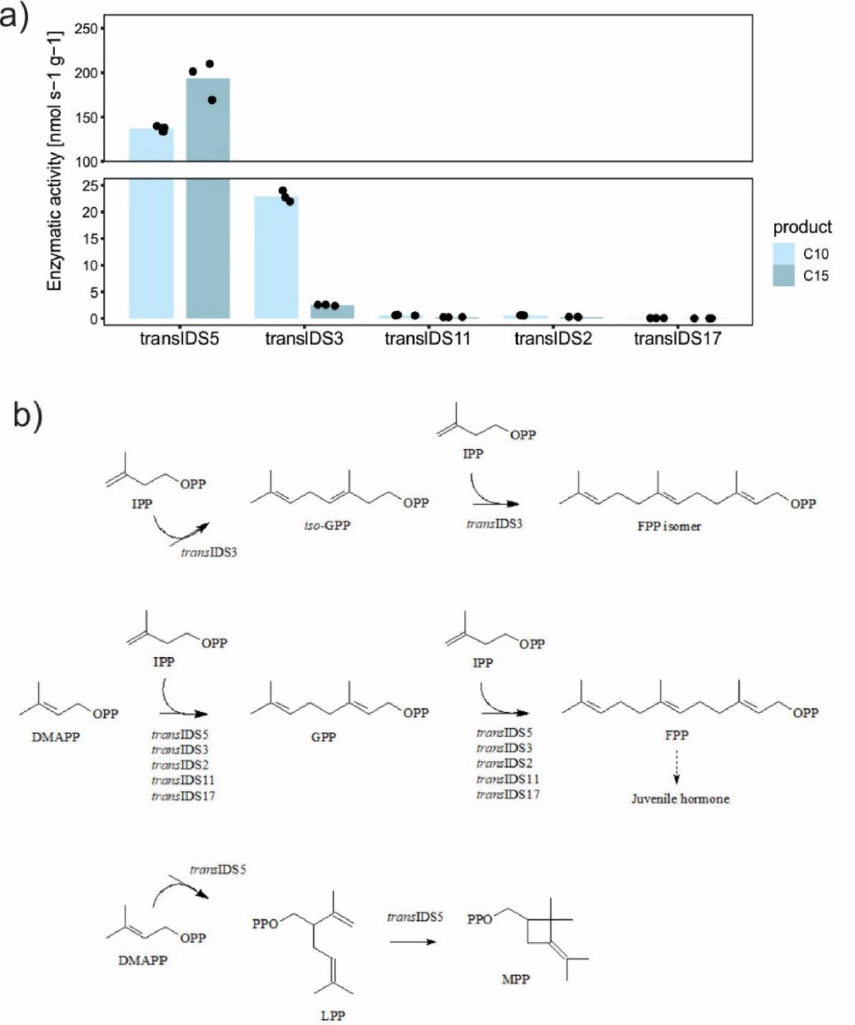
*In vitro* activity of *P. citri* IDS sequences expressed in *E. coli*. Measured enzymatic activity for production of C10 and C15 prenyl diphosphates (a) and schematic representation of confirmed coupling reactions performed by *P. citri trans*-IDS candidates (b). *Trans*IDS5, *trans*ID11, *trans*IDS2, and *trans*IDS17 produced C10 geranyl diphosphate (GPP) and C15 farnesyl pyrophosphate (FPP); *trans*IDS3 produced C10 GPP and *cis*-isogeranyl diphosphate and two C15 FPP isomers. (a) Activity for production of C10 prenyl diphosphates (light blue bars) and activity for production of C15 prenyl diphosphates (dark blue bars). Height of the bars represents the mean of three separate measurements, plotted as block dots. Raw data is available in Table S5. Note the y-axis break due to much higher activity of *trans*IDS5. We did not detect any C10 or C15 products with other tested candidates (Table 1). In (b), dimethylallyl diphosphate (DMAPP) can be joined to its isomer isopentenyl diphosphate (IPP) to form regular terpenes geranyl and iso-geranyl diphosphates (GPP and iso-GPP) and isomers of farnesyl diphosphate (FPP). Alternatively, two DMAPP units can be assembled into irregular lavandulyl diphosphate (LPP) and maconelliyl diphosphate (MPP). FPP is a precursor for juvenile hormone biosynthesis as denoted by the dashed arrow.

Moreover, candidate *trans*IDS5 also had a low level of irregular IDS activity, if supplied with DMAPP only (Figure 3b). The enzyme produced two compounds, which were identified as lavandulyl (LPP) and maconelliyl diphosphate (MPP), based on GC-MS spectra of their dephosphorylated derivatives (Figure S20).

To assess the activities of candidate proteins *in vivo* in the eukaryotic cell environment, the candidate IDS sequences were transiently expressed in the plant heterologous expression model *Nicotiana benthamiana*. HPLC-MS analysis of leaf extracts did not reveal irregular terpene diphosphates. Similarly, no production of volatile organic compounds derived from irregular monoterpenoids was detected by GC-MS in agroinfiltrated *N. benthamiana* leaves. Regular activity of candidate proteins could not be assessed in this system due to the presence of native plant regular monoterpenes.

### Mutagenesis of *trans*IDS5 active site revealed functional importance of active site residues for its enzymatic activity

We further set to identify the amino acid residues important for the irregular coupling activity of *trans*IDS5 by performing mutagenesis. It was suggested that replacement of the first aspartate (D) in SARM with asparagine (N) played an important role in the evolution of irregular coupling activity (Liu et al., 2012). However, in *trans*IDS5, analogous mutation (D308N) caused a complete loss of irregular activity. The same was observed after the exchange of two other aspartate residues in SARM (D309N and D312N), and the first aspartate in FARM (D166N).

Moreover, the regular activity of aspartate mutants was affected to different degrees (Figure 4a). The regular activity of D166N mutant was significantly diminished, although the ratio between C10 GPP and C15 FPP was similar to the wild-type (wt) *trans*IDS5 (Figure 4a). D308N and D312N substitutions had impact on the regular product length. Both mutants performed the initial coupling of IPP and DMAPP to C10 chains at levels comparable to the wt enzyme, but the addition of the second IPP to form the C15 chain was hampered compared to the wt (Figure 4a). The regular activity was completely abolished after the D309N exchange (Figure 4a).

**Figure 4:**
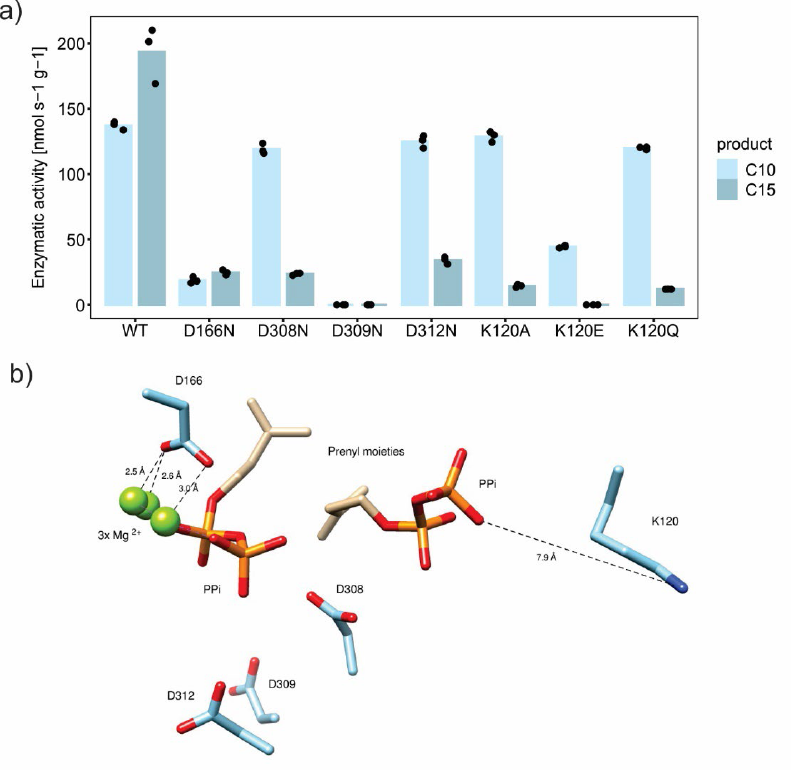
Mutagenesis of *trans*IDS5 active site residues. (a) Activity of *trans*IDS5 and its mutants expressed in *E. coli.* Activity for production of C10 geranyl diphosphate (GPP, light blue bars) and activity for production of C15 farnesyl diphosphate (FPP, dark blue bars) are plotted. Height of the bars represents the mean of three separate measurements, plotted as black dots. Raw data is available in Table S5. (b) Mutated aminoacids and their position in the active site, based on a computational model of active centre of *trans*IDS5 with two bound substrate molecules showing the aspartate of the first (D166) and the second (D308, D309, D312) aspartate-rich motifs and lysine 120 (K120). Oxygen atoms in aspartate side chains and in pyrophosphate moieties (PPi) of the substrates are shown in red; the nitrogen atom of lysine side chain is highlighted in blue.

To find additional residues important for the catalytic activity of *trans*IDS5, we build a computational model of its 3D structure (Figure 4b). We identified a positively charged lysine (K) residue in position 120 located at a comparatively short distance from the negatively charged diphosphate moiety of the substrate (Figure 4b). We exchanged this residue with alanine (K120A), polar uncharged glutamine (K120Q), and negatively charged glutamate (K120E). All three substitutions resulted in the loss of irregular coupling activity, as well as in the diminished elongation of GPP (C10) to FPP (C15). As expected, the K120E exchange, introducing negative charge in the active site, had the most inhibiting effect, precluding FPP formation entirely (Figure 4a).

### Heteromeric associations did not change IDS activity

In most cases, *trans*- and *cis*-IDSs are active in the form of homodimers (Gao et al., 2012). However, some IDSs form heterodimers with non-catalytic subunits, which modulate or stimulate their activity (Burke et al., 1999; Tholl et al., 2004; Qu et al., 2015; Yin et al., 2017; Zhou et al., 2017; Emi et al., 2019; Dong et al., 2023). To determine if irregular coupling might depend on the presence of other subunits, we tested *P. citri* IDS candidates in combinations with potential interactors (Table S6). In the *P. citri* genome, we identified homologs of the DPPS and DHPPS regulatory subunits (*trans*IDS4 and *cis*IDS9, respectively) and both were combined with all IDS-like candidates in irregular activity assays (Table S6). In addition, we examined two *P. citri* non-IDS-like proteins for their IDS-modifying activities: the protein product of g23689.t1, annotated as a pheromone-binding protein (PBP) and highly expressed in virgin females (Table S3b), and g36424.t1, identified as a *P. citri* homolog of isopentenyl diphosphate isomerase (IDI), an enzyme catalysing the reversible isomerisation of IPP to DMAPP (Table S6). Addition of *P. citri*-IDI to the reaction with IDS candidates supported formation of regular products, if either IPP or DMAPP only were added to the reaction. However, none of the tested putative regulatory subunits altered or spurred irregular coupling activity of IDS candidates.

We also tested the hypothesis that heterodimeric complexes consisting of two IDS subunits are required for irregular coupling. Activity assays with DMAPP as the only substrate were carried out with pairwise combinations of *P. citri* IDS candidate proteins with each other as well as with IDS proteins forming irregular prenyl diphosphates (Table S6). The latter allows for detection of terpene synthase (TPS) activities of *P. citri* IDS candidates, converting the prenyl diphosphate substrates into corresponding prenyl alcohols. Combined assays did not reveal any changes to the enzymatic activities of IDS candidates, which also did not demonstrate TPS activity on the tested irregular prenyl diphosphates.

## Discussion

Sustainable insect pest management, supported by state-of-the-art knowledge and technology, is an integral part of sustainable food production in a changing climate, in which the geographical ranges of many insect species are expected to widen (Schneider et al., 2022). However, the use of biotechnology and genetic solutions depend on detailed information about gene function. In this work, we present the results of a comprehensive search for candidate genes coding for isopentenyl diphosphate synthase (IDS) activity from the citrus mealybug, *P. citri*. The identification of these genes has the potential to enable either biological production of *P. citri* sex pheromones or gene silencing approaches, both of which provide novel options for pest management (reviewed in Mateos Fernández et al., 2022).

Functional genomics of many insect species is hindered by the lack of well-annotated, high-quality genome sequences (Li et al., 2019; Hotaling et al., 2021). To improve the available genetic resources for *P. citri,* we complemented the existing genome data with short- and long-read transcriptome data (Figure 1). The transcriptomic dataset enabled us to predict and amplify alternative transcript sequences (Figure 2, Figure S3). In addition, we found a novel coding sequence (*trans*IDS5). Using all resources, we were able to identify 18 candidate *P. citri* IDS sequences. Among these were putative DHPPS, FPPS, FPPS-like, GGPPS, DPPS and DPPS-like coding sequences, as well as an ubiquitin thioesterase-like sequence with a D-rich motif (Table 1).

However, inferring function from sequence analyses alone is inconclusive for IDSs, as even small changes can alter substrate and chain-length specificity (Wallrapp et al., 2013). To test the activity of IDS candidates, we focused on the detection of C10 and C15 coupling products. We, therefore, did not confirm if the predicted DHPPS, GGPPS and DPPS had longer chain-producing activities. *Trans*IDS16, a predicted GGPPS did not produce any C10 or C15 products. Similarly, neither of the two tested *cis*-IDS-like proteins produced detectable levels of regular or irregular C10 or C15 backbones, as both were predicted to be long-chain IDSs, namely DHPPS (*cis*IDS1) and UPPS (*cis*IDS8) (Table 1). Interestingly, *trans*IDS2, a predicted long-chain IDS (DPPS catalytic subunit), produced low amounts of FPP and GPP. This might be attributed to functional promiscuity *in vitro*, producing polyprenyl chains of different lengths, as reported for other IDS enzymes (Lackus et al., 2019). Additionally, *trans*IDS17, which was not homologous to IDS sequences but rather to otubain-like ubiquitin thioesterases, was able to produce regular C10 and C15 backbones, albeit at the lowest rate of all candidates. It remains unknown if otubain-like sequences from other mealybug species have IDS activity, and whether this activity has a biological function.

The two putative FPPS sequences (*trans*IDS3 and *trans*IDS5) both displayed regular coupling activity resulting in C10 and C15 units. Our enzymatic activity and gene expression results indicate that *trans*IDS5 might be the major source of C15 terpenes in *P. citri*, while the C10 could be mainly composed of *trans*IDS5 and possibly also *trans*IDS3 products. The highest specific activity was measured for *trans*IDS5, also producing low amounts of irregular prenyl diphosphates, LPP and MPP (Figure 3b), which have not been reported in *P. citri*. The 2-methylbutanoated versions of both compounds have been identified as the sex pheromone of *Maconellicoccus hirsutus*, the pink hibiscus mealybug (Zhang et al., 2004). Esterified lavandulol compounds act as sex pheromones also in some other *Planococcus*, *Pseudococcus* and *Dysmicoccus* species (Zou and Millar, 2015). However, the irregular coupling activity of *trans*IDS5 only takes place in the absence of IPP and might, therefore, be unlikely to occur *in vivo* where the presence of IPP is expected. Although we did not find conclusive evidence for the role *trans*IDS5 in the biosynthesis of the *P. citri* sex pheromone, we propose that it is the main source of regular sesquiterpenes in *P. citri*, possibly including the juvenile hormone (Figure 3b). Using structure-informed mutagenesis, we additionally demonstrated the importance of K120, D166, D308, D309, and D312 for the irregular coupling mechanism observed for *trans*IDS5 *in vitro* (Figure 4a).

Terpenoids are known for their extensive evolutionary divergence, with many instances of lineage-specific pathways and, therefore, metabolites. Despite the occurrence of many common and unique terpenoids in animals, terpenoid diversification is better understood in plants and microbes (Beran et al., 2019; Tholl et al., 2023). In plants, terpenoid diversity is mainly determined by the activity of TPSs, which have high evolutionary divergence and functional plasticity (Karunanithi and Zerbe, 2019). However, sequences with homology to plant and microbial TPSs have not been identified in insect genomes. Instead, numerous instances of IDS gene-family expansion have been observed, with frequent gene duplications resulting in functional redundancy or neofunctionalization (Gilg et al., 2009; Beran et al., 2016; Lancaster et al., 2018, 2019; Darragh et al., 2021; Rebholz et al., 2023). Among the *P. citri* IDS sequences, we observed expansions of FPPS-like sequences and DHPPS-like sequences (Figure S9, Figure S12). The former extends the previously reported FPPS-like diversification within the Coccomorpha (Rebholz et al., 2023). Three FPPS-like sequences were successfully cloned and tested (*trans*IDS10, *trans*IDS11, and *trans*IDS12), however, only one candidate (*trans*IDS11) exhibited IDS activity, producing low amounts of GPP and FPP. Interestingly, while we were not able to confirm GPPS or FPPS activity for *trans*IDS12, its expression was observed to be strongly upregulated in virgin compared to mated *P. citri* females, which was also true for *trans*IDS9, *trans*IDS10, and *trans*IDS11 (Table 1). This differential expression suggests a possible role in mating or reproduction and needs further investigation.

Despite detecting two irregular prenyl diphosphate products, none of the IDS-like candidates that we tested displayed coupling activity leading to *cis*-planococcyl diphosphate under our experimental conditions. *In vitro* functional characterization of IDS enzymes can be unreliable, as activity can depend on many factors, including the formation of heteromers (Zhang and Li, 2013). To that end, we tested for modulating activity of several potential interactors but did not detect any changes in the activity of *P. citri* IDS candidates.

The model of irregular terpenoid biosynthesis via a genome-encoded IDS has been challenged. Alternative hypotheses include the possibility that insects sequester plant-derived compounds or utilise products of their endosymbionts’ metabolism. Many studies reveal a straightforward strategy of modification and accumulation of plant defence compounds (Beran and Petschenka, 2022), but, in some cases, the derivatives of plant secondary metabolites are also used in sexual communication. For example, fragments of plant pyrrolizidine alkaloids and phenylpropanoid metabolites are reported to make danaine butterfly males more attractive to the co-specific females (Nishida et al., 1991). However, the presence of irregular monoterpene structures has not been reported for known *P. citri* host plants which makes the sequestration hypothesis improbable. Like other insects feeding on nutritionally unbalanced plant sap, mealybugs rely on obligate endosymbiotic bacteria as a source of amino acids and vitamins (Husnik et al., 2013). *P. citri* is a part of a nested tripartite endosymbiotic system, where the insect harbours the Betaproteobacteria *Candidatus* Tremblaya princeps, which contains the Gammaproteobacteria *Candidatus* Moranella endobia (McCutcheon and Dohlen, 2011; López-Madrigal et al., 2013). Although we were unable to identify any short-chain IDS-coding sequences within the genomes of *P. citri* endosymbionts, we cloned and tested one IDS sequence, a putative UPPS, from *Candidatus* Moranella endobia (*cis*IDS8). However, this did not produce any regular or irregular short prenyl chains. Moreover, genetic crosses between *P. citri* and *Planococcus minor* - the latter producing a branched lavandulol-type monoterpene - indicate that the enzyme responsible for the formation of the irregular terpene skeleton is encoded in the insect nuclear genome and is present at a single locus (Tabata, 2022). The structure of the sex pheromone is also not inherited maternally (Tabata, 2022), as would be expected if synthesis requires the transfer of egg-transmitted endosymbionts.

Another possibility is that the mealybug sex pheromones are biosynthesised via a different catalytic mechanism, executed by a different, and as yet unidentified, class of enzymes. Novel terpene synthases have been described, expanding on the canonical models of terpene biosynthesis (Rudolf and Chang, 2020). A recent example is the elucidation of the iridoid pheromone found in the pea aphid, *Acyrthosiphon pisum*. Biosynthesis of this molecule proceeds through the same intermediates as in plants, however, it was shown to involve a set of enzymes which lack homology to the plant counterparts (Köllner et al., 2022). The insect enzymes were identified by transcriptomic analysis of the female hind legs, the site of pheromone biosynthesis (Köllner et al., 2022). Targeted transcriptome analyses of pheromone glands were also useful for determining the genes related to pheromone metabolism and transport in Lepidoptera species (Vogel et al., 2010; Gu et al., 2013; Nuo et al., 2021; Yao et al., 2021). At present, however, such approaches are not possible in *P. citri* as the site of the pheromone gland remains unknown.

The identification of the *P. citri* IDS gene family provides an important foundation for deciphering terpenoid biosynthesis in this species as well as other mealybugs. The candidate FPPS and FPPS-like sequences with confirmed regular and irregular activity might also present a starting point for protein engineering; mutations have been shown to increase or even spur irregular coupling activity in IDSs (Chan et al., 2017; Gerasymenko et al., 2022). Diversification of IDS sequences in Coccoidea is an intriguing genomic enigma for future investigation, especially in relation to their capacity for species-specific irregular monoterpene synthesis, a unique feature in this insect superfamily, which still remains elusive.

## Methods

### Insect rearing and sampling

The stock colony of *P. citri* was established in the insectary facilities at the Universitat Politècnica de València (UPV, Valencia, Spain) using specimens from Servei de Sanitat Vegetal of Generalitat Valenciana (GVA, Valencia, Spain). Mealybugs were reared on organic green lemons, maintained in a rearing chamber, in the dark at 24 ± 2 °C, with 40−60 % relative humidity. Mated females were allowed to oviposit and, when ovisacs were laid, they were gently transferred with an entomological needle to new lemons. To obtain samples of virgin females, some lemons were separated from the main colony and were visually inspected every 3−4 days for the presence of male cocoons, which were manually removed with an entomological needle to leave only virgin females. Mated females were sampled on the lemons from the main stock colony after checking for the presence of ovisacs. The production of sex pheromone was confirmed by collection of emitted volatiles from a group of 20−30 virgin females by solid phase microextraction technique (SPME) (Vacas et al., 2017). The identity of the sex pheromone was unequivocally confirmed by full coincidence of its mass spectra and retention time with an analytical standard sample synthesised according to the method described by Kukovinets et al., 2006.

### RNA isolation

RNA was isolated from approximately 150 (cca. 100 mg) pooled virgin and mated *P. citri* females separately, using TRIzol Reagent (Thermo Fisher Scientific, Waltham, MA, USA) according to the manufacturer’s instructions. RNA was isolated from four biological replicates, resulting in eight RNA samples, four from pools of virgin females and four from pools of mated females. RNA quality was assessed using the Quant-iT™ RNA Assay Kit (Thermo Fisher Scientific, Waltham, MA, USA).

### Short read Illumina RNA-Seq

Approximately 1 µg of isolated RNA from all eight RNA samples was purified to extract polyadenylated mRNA using biotin beads, fragmented, and primed with random hexamers to produce the first cDNA strand followed by second-strand synthesis to produce double-stranded cDNA. Libraries were constructed at the Earlham Institute, UK, on a Sciclone G3 NGSx workstation (PerkinElmer, Waltham, MA, USA) using the TruSeq RNA protocol v2 (Illumina Part # 15026495 Rev.F) and sequenced on two lanes of a HiSeq 4000 (Illumina, San Diego, CA, USA) generating 150 bp negative strand-specific paired-end reads. Illumina reads were subjected to adapter trimming, read quality filtering, mapping and read summarization (counting only uniquely mapped paired reads) in CLC Genomics Workbench 10.0.1 (Qiagen, Hilden, Germany), with mapping parameters: mismatch cost 2, insertion cost 3, deletion cost 3, length fraction 0.9, similarity fraction 0.9. Reads were mapped to the *Pcitri*.v1 genome, downloaded from the MealyBugBase (https://ensembl.mealybug.org/index.html). Differential expression analysis was performed with R packages edgeR and limma (Law et al., 2018), contrasting samples from virgin and mated *P. citri* females.

### Long-read Iso-Seq transcriptome

Approximately 2 µg of RNA isolated from pooled virgin *P. citri* females was sent for PacBio SMRT sequencing using the Iso-Seq protocol (National Genomics Infrastructure, Sweden). High and low quality full-length transcript isoforms were mapped to the *Pcitri*.v1 genome using minimap2 (Li, 2018). Based on the genome mapping, the isoforms were collapsed and representative isoforms were filtered to exclude isoforms with 5’ artefacts, all using scripts from cDNA_cupcake Github repository (Tseng, 2022).

### *De novo* short-read transcriptome assembly

Trimmed paired and orphan Illumina RNA-Seq reads were subjected to *de novo* transcriptome assembly using rnaSPAdes v3.13.0 (Bushmanova et al., 2019). Two assemblies were created, first, using the virgin and mated female *P. citri* paired-end short reads (2×150 nt) obtained in this study, and second, combining them with paired-end reads of seven samples of *P. citri,* kindly provided by Dr. Laura Ross at the University of Edinburgh (2×75 nt and 2×50 nt, see Data Availability Statement). For the first assembly, default rnaSPAdes parameters were used, whereas for the second assembly, adapter trimming and sequencing artefact removal was additionally performed using BBTools’ bbduk script (Bushnell, 2022), followed by SPAdes’ BayesHammer (Nikolenko et al., 2013) and BBTools’ ecco, ecc (6 passes) and tadpole read error correction scripts. Assembly was done using k-mer lengths 29 and 49.

### Consolidation of sequence resources, annotation and quality control

The two Illumina short read assemblies (Figure 1a and b) were combined into a single set (Figure 1c) by removing identical sequences with CD-HIT-EST v6.4 (Fu et al., 2012) using 100% sequence identity threshold. In a successive run, the combined short read transcriptome and the Iso-Seq transcriptome (combining mapped and unmapped representative isoforms, Figure 1d) were consolidated into a single set (Figure 1e), again using CD-HIT-EST at 100% sequence identity threshold. The final consolidated transcriptome dataset as well as both short- and long-read datasets were quality checked using rnaQUAST (Bushmanova et al., 2016) and their completeness was assessed using BUSCO v5.4.3 (Manni et al., 2021) with insecta_odb10 lineage dataset. The consolidated transcriptome set was mapped to the *Pcitri*.v1 genome using STARlong v2.7.5c (Dobin et al., 2013) and mapped transcripts were matched with the gene models and scaffolds using matchAnnot (Skelly, 2015; version 20150611.02). Transcripts were also subjected to ORF prediction and translation using Transdecoder v5.5.0 (Haas, 2023), followed by the annotation of the predicted protein sequences with PFAM and InterPro (IPR) IDs using InterProScan v5.56-89.0 (Jones et al., 2014). The longest ORFs were further used for taxonomic classification using mmseq2 taxonomy tool (Mirdita et al., 2021), querying against the non-redundant NCBI protein database (downloaded on 19.07.2022). Results of taxonomic classification were visualised with Pavian (Breitwieser and Salzberg, 2020). Illumina short reads were mapped back to both *de novo* assemblies as well as IsoSeq transcriptome using STAR v2.7.5c (Dobin et al., 2013) with default parameters. Short read mapping counts for all three datasets were used for differential transcript expression analysis based on R packages edgeR and limma (Law et al., 2018).

### Candidate selection

We queried the IPR annotations of *Pcitri*.v1 gene models and our consolidated transcriptome dataset with IDS-related IPR IDs, namely IPR001441 (Decaprenyl diphosphate synthase-like family), IPR036424 (Decaprenyl diphosphate synthase-like superfamily) and IPR000092 (Polyprenyl synthetase family), to identify homologs of *cis*- and *trans*-prenyltransferases. Besides using the IPR search to determine *trans*-IDS homologs, we also conducted a motif search using MAST v5.0.1 in MEME Suite (Bailey et al., 2015). Twenty-one sequences of farnesyl pyrophosphate synthases (FPPS) from different organisms were used as an input for MEME. The identified motifs were used as a query against the *Pcitri*.v1 proteins (Table S7). Additionally, we also searched for *P. citri* isopentenyl diphosphate isomerase (IDI) homologs by querying the *P. citri*.v1 protein sequences with *Nicotiana tabacum* IDI sequence (GenBank accession NP_001313140) using blastp (Altschul et al., 1990).

All extracted sequences with homology to *cis*- and *trans*-IDSs longer than 100 amino acids were subjected to CD-HIT with 100% sequence identity threshold to remove duplicates. They were further grouped into clusters, using CD-HIT along with multiple sequence alignments with MUSCLE (Edgar, 2004), and the sequence representing the most probable full-length coding sequence was selected within each cluster.

For the phylogenetic analyses of candidate sequences, we first searched for similar sequences using blastp against the non-redundant protein bases or tblastn against the transcriptome shotgun assembly sequences (TSA). Additionally, we searched for homologs from *Planococcus ficus* and *Pseudococcus longispinus* genome assemblies in the MealyBugBase (https://ensembl.mealybug.org/index.html), and from *Maconellicoccus hirsutus* transcriptomic resource (Kohli et al., 2021). Amino acid sequences were aligned using the MUSCLE algorithm (Edgar, 2004) and used to construct maximum-likelihood phylogenetic trees with LG model (Le and Gascuel, 2008) tested with the bootstrap method at 1000 replicates in MEGA-X, version 10.0.5 (Kumar et al., 2018).

### Candidate sequence amplification and expression in *E. coli*

The candidate sequences were amplified from the cDNA prepared with oligoT primer (T15NNN) from total RNA isolated from virgin *P. citri* females using primers listed in Table S8 and cloned into pMAL-c5X expression vector (New England Biolabs, Ipswich, MA, USA) using *Nco*I/*Bam*HI sites. The proteins were expressed in BL21(DE3)pLysS *E. coli* cells. The expression was induced by 0.5 mM IPTG and carried out at 30 °C overnight. The proteins were purified on the amylose resin (New England Biolabs, Ipswich, MA, USA) and stored at −20 °C in 25 % glycerol. The protein concentration was determined by the method of Bradford (Bradford, 1976).

### Enzyme activity assays

Isopentenyl, dimethylallyl, and geranyl diphosphate (IPP, DMAPP, and GPP) were purchased as tri-ammonium salts from Echelon Biosciences (Salt Lake City, UT, USA). The compounds were dissolved in the mixture of methanol:water (7:3) at the concentration of 1 mg/mL and stored at −20 °C. The activity assays were carried out using 10–180 μg of protein in a total volume of 100 μL in 35 mM HEPES buffer, pH 7.4, containing 10 mM MgCl_2_ and 5 mM β-mercaptoethanol at 30 °C for 30 or 60 min. Regular coupling activity was determined using IPP and DMAPP as substrates, both at 100 μM, while the assays for irregular activity contained 200 μM of only DMAPP. The reaction was stopped by adding 100 μL of chloroform, and the proteins were precipitated by vigorous shaking followed by centrifugation (10 min at 16,000 g). The water phase (injection volume 5 μL) was used for LC-MS analysis performed on an 1260 Infinity HPLC system coupled to a G6120B quadrupole mass spectrometry detector (Agilent Technologies, Santa Clara, CA, USA). The separation was carried out on a Poroshell 120 EC-C18 (3.0 x 50 mm, 2.7 μm) column (Agilent Technologies, Santa Clara, CA, USA) using the method described in (Nagel et al., 2012). The detection of C10 and C15 prenyl diphosphates was performed in negative single ion mode; m/z 313 and 381, respectively. The quantity of formed products was calculated based on peak areas using a calibration line built for GPP.

For analysis of regular terpene alcohols, the reactions were carried out with 70–330 μg of protein in a total volume of 400 μL in 35 mM HEPES buffer, pH 7.4, containing 10 mM MgCl_2_, 5 mM β-mercaptoethanol, and substrates (1 mM IPP and 1 mM DMAPP) at 28 °C overnight. For dephosphorylation of prenyl diphosphates, the reaction mixture was supplemented with 80 μL of 0.5 M glycine-NaOH buffer (pH 10.5) containing 5 mM ZnSO_4_; 80 units of calf intestinal alkaline phosphatase (Promega, Madison, WI, USA) were added and the mixture was incubated at 37 °C for 1 h. The alcohols were extracted 3 times with 400 μL of methyl tert-butyl ether (MTBE); the organic phase was evaporated to approximately 150 μL under compressed air flow. GC-MS analysis was carried out using DB-5ms column (30 m x 0.250 mm x 0.25 μm; Agilent Technologies, Santa Clara, CA, USA) with the following temperature gradient: 50 °C, from 50 °C to 120 °C at 7 °C per min, 120 °C to 320 °C at 35 °C per min, hold 5 min. Carrier gas (H_2_) flow was set at 3 mL/min. Injection volume was 1 μL (PTV splitless injection, initial injector temperature 60 °C, final injector temperature 250 °C, injector heating rate 50 °C/sec).

For identification of irregular products of *trans*IDS5, 1 mg of protein was incubated with 2 mg of DMAPP tri-ammonium salt in 1.6 mL of 35 mM HEPES buffer, pH 7.4, containing 10 mM MgCl_2_, and 5 mM β-mercaptoethanol at 28 °C overnight. The products were dephosphorylated and extracted as described above, and separated on ALUGRAM^®^ Xtra SIL G/UV_254_ TLC plates (0.20 mm silica gel layer, Macharey-Nagel, Düren, Germany) using hexane:acetone (4:1) mixture. Individual substances were extracted from silica gel with MTBE and analysed by GC-MS using Cyclosil-B capillary column (30 m x 0.250 mm x 0.25 μm; Agilent Technologies, Santa Clara, CA, USA) with the following temperature gradient: 50 °C for 3 min → from 50 °C to 120 °C at 2 °C per min → 120 °C to 230 °C at 10 °C per min. Helium was applied as the carrier gas at a flow rate of 1 mL/min. Injection volume was 1 μL (1:100 split).

### Candidate expression in *Nicotiana benthamiana*

Genetic constructs for candidate expression in plants were assembled with the Golden Braid (GB) cloning system (Sarrion-Perdigones et al., 2011). Transcriptional units in alpha-level GB vectors included IDS coding sequences fused with a C-terminal His-tag under control of the 35S CaMV promoter. Each candidate gene was tested in its native form and with the addition of a modified chloroplast transit peptide (cTP) of ribulose bisphosphate carboxylase small subunit from *Nicotiana sylvestris* (GenBank accession number XP_009794476.1). If a different (e.g. mitochondrial) localization signal was predicted, it was removed before adding the cTP. Transient expression in *N. benthamiana* leaves was carried out as described earlier (Gerasymenko et al., 2019). For detection of irregular IDS activity, the plant material (200 mg) was frozen in liquid nitrogen, ground and extracted with 80 % methanol (400 μL) by 30 min incubation in the ice ultrasound bath. Supernatants after two centrifugation steps (10 min at 16,000 g) were analysed using 1260 Infinity HPLC system coupled to G6120B quadrupole mass spectrometry detector (Agilent Technologies, Santa Clara, CA, USA). The separation was carried out on Zorbax Extend-C18 (4.6 x 150 mm, 3.5 μm) column (Agilent Technologies, Santa Clara, CA, USA) with mobile phase consisting of 5 mM ammonium bicarbonate in water as solvent A and acetonitrile as solvent B applying the following gradient (% B): starting at 0, 0 to 8 within 2 min, holding 8 for 15 min, 8 to 64 within 3 min, 64 to 100 within 2 min, holding 100 for 4 min, 100 to 0 within 1 min, re-equilibrating at 0 for 12 min. The detection of C10 and C15 prenyl diphosphates was performed in negative single ion mode; m/z 313 and 381, respectively. For detection of volatile compounds, samples from agroinfiltrated leaves were collected 5 dpi, snap-frozen in liquid nitrogen and analysed by GC-MS as described previously (Mateos-Fernández et al., 2021).

### Computational model and site-directed mutagenesis of *trans*IDS5

The 3-D model of *trans*IDS5 was built on the SWISS-MODEL homology-modelling server (Waterhouse et al., 2018) using the crystal structure of *Saccharomyces cerevisiae* geranylgeranyl pyrophosphate synthase in complex with magnesium and IPP (PDB DOI: 10.2210/pdb2E8U/pdb) as a template. For structure visualisation and analysis, UCSF Chimera v.1.14 was applied (Pettersen et al., 2004). Mutations were introduced using the Q5-SDM kit (New England Biolabs) with primers listed in Table S8.

## Acknowledgements and Funding

All authors gratefully acknowledge the European Research Area Cofund Action ‘ERACoBioTech’ for the support of the project SUSPHIRE (Sustainable Production of Pheromones for Insect Pest Control in Agriculture), which received funding from the Horizon 2020 research and innovation program under grant agreement No. 722361. M.J., M.P., K.G., and Š.B. acknowledge the financial support from the Slovenian Ministry of Education, Science and Sport, as well Slovenian Research Agency (grant No. P4-0165). I.M.G., E.H., and H.W. acknowledge the support by German Federal Ministry of Education and Research (BMBF), grant number 031B060. K.K. and N.P. acknowledge the support of the UK Biotechnology and Biological Sciences Research Council (BBSRC) Core Strategic Program Grant to the Earlham Institute (Genomes to Food Security; BB/CSP1720/1) and grant BB/R021554/1. S.G. and D.O. acknowledge grant PLEC2021-008020 (PHEROPLUS) by the Spanish Ministry of Science and Innovation, the Next Generation EU initiative and the Spanish Agencia Estatal de Investigación (AEI). S.G. acknowledges a postdoctoral grant (CIAPOS/2021/316) from the Generalitat Valenciana and the Fondo Social Europeo. Illumina sequencing was delivered via the BBSRC National Capability in Genomics and Single Cell Analysis (BBS/E/T/000PR9816) at the Earlham Institute by members of the Genomics Pipelines Group. I.M.G. is grateful to Prof. Dr. Michael Heethoff from the Technical University of Darmstadt and to Prof. Dr. Richard Dehn, Laura Kleditzsch, and Miriam Peters from the University of Applied Sciences in Darmstadt for the possibility to carry out GC-MS measurements. We would also like to thank Dr Laura Ross and her group from the University of Edinburgh for kindly providing their *P. citri* Illumina datasets as well as Dr Rosario Gil Garcia from the University of Valencia for providing insights into the *P. citri* endosymbionts and their genomes.

## Data and code availabilitys

Raw Illumina reads from *P. citri* virgin and mated females, along with results of the differential expression analysis (as given in Table S3b as well), were deposited at GEO under accession GSE179660. Raw Iso-Seq reads were deposited at SRA under accession SRR15093694. *P. citri* short reads provided by the University of Edinburgh and used in the second *de novo* transcriptome assembly are deposited at SRA under accession numbers SRR11260462, SRR11260463, SRR11260468, SRR11260469, SRR11260470, and SRR11260471. *De novo* transcriptome assembly FASTA file is available at FAIRDOMHub (https://fairdomhub.org/data_files/6316/content_blobs/17124/download), together with an annotation file (https://fairdomhub.org/data_files/6317/content_blobs/17125/download), also included in the Supplemental Files (Table S3a).

## Supplemental Files

**Table S1:** Quality assessment of short- and long-read transcriptome data.

**Table S2:** Percent of short reads mapping to transcriptome sequences.

**Table S3:** Differential expression of *P. citri* genes between virgin and mated females.

**Table S4:** Selected IDS sequences.

**Table S5:** IDS activity measurements of *P. citri* candidates.

**Table S6:** Combined activity assays performed for detection of irregular IDS and terpene synthase activities.

**Table S7:** Input and output of the MEME motif search for *P. citri* sequences containing *trans*-IDS motifs.

**Table S8:** Primers used for amplification of IDS sequences from *P. citri* cDNA and for introducing mutations into *trans*IDS5 sequence.

**Figure S1:** BUSCO assessment of transcriptome completeness.

**Figure S2:** Taxonomic classification of the *P. citri* transcriptome dataset.

**Figure S3:** Alignments of sequence resources for *trans*IDSl0 and *trans*IDS13.

**Figure S4:** Phylogenetic tree of putative DPPS subunit 1 sequences from selected species with *trans*IDS2 and *trans*IDS7.

**Figure S5:** Phylogenetic tree of putative FPPS sequences from selected species with *trans*IDS3, *trans*IDS5, and *trans*IDS6.

**Figure S6:** Phylogenetic tree of putative DHPPS catalytic subunit sequences from selected species with *cis*IDS1, and *cis*IDS4.

**Figure S7:** Phylogenetic tree of putative DHPPS regulatory subunit sequences from selected species with *cis*IDS6, and *cis*IDS9.

**Figure S8:** Phylogenetic tree of putative DHPPS regulatory subunit sequences from selected species with *cis*IDS6, and *cis*IDS9.

**Figure S9:** Phylogenetic tree of putative FPPS and FPPS-like sequences from Coccoidea species.

**Figure S10:** Phylogenetic tree of putative GGPPS sequences from selected species with *trans*IDS16.

**Figure S11:** Phylogenetic tree of putative otubain-like sequences from selected species with *trans*IDS17.

**Figure S12:** Phylogenetic tree of putative DHPPS and DHPPS-like sequences from selected Coccoidea species.

**Figure S13:** Identification of regular monoterpenes synthesised by *trans*-IDS enzymes from *P. citri*.

**Figure S14:** EI mass spectra of geraniol generated by dephosphorylation of standard GPP (1A) and products of *trans*IDS5 (1B) and *trans*IDS3 (1C).

**Figure S15:** EI mass spectra of geraniol generated by dephosphorylation of standard GPP (1A) and products of *trans*IDS5 (1B) and *trans*IDS3 (1C).

**Figure S16:** Identification of regular sesquiterpenes synthesised by *trans*-IDS enzymes from *P. citri*.

**Figure S17:** EI mass spectra of *trans*-farnesol generated by dephosphorylation of products of FPPS from *Tanacetum cinerariifolium* known to produce *trans*-farnesyl diphosphate (4A) and *trans*IDS5 (4B).

**Figure S18:** EI mass spectra of dephosphorylated products of *trans*IDS3, *trans*-farnesol (4C) and farnesol isomer (5).

**Figure S19:** Identification of regular monoterpene diphosphates synthesised by *trans*-IDS enzymes from *P. citri*.

**Figure S20:** Identification of irregular monoterpenes synthesised by *trans*IDS5.

