## Supplemental Figures and Tables for "Identification and characterisation of *Planococcus citri cis*- and *trans*-isoprenyl diphosphate synthase genes, supported by short- and long-read transcriptome data"

[5] Instituto Agroforestal del Mediterráneo-CEQA, Universitat Politècnica de València. Camino de Vera s/n, Valencia, Spain

[6] Engineering Biology, Earlham Institute, Norwich Research Park, Norwich, Norfolk, NR4 7UZ, UK

[7] Institute for Plant Molecular and Cell Biology (IBMCP), Consejo Superior de Investigaciones Científicas (CSIC) - Universitat Politècnica de València (UPV), Valencia, Spain

[8] Department of Organic Chemistry, University of Valencia, Burjassot, València, Spain

[#] These authors contributed equally to the work.

*Coresspondence to

**Table S1: Quality assessment of short- and long-read transcriptome data.** (a) Combined rnaQUAST outputs for the consolidated transcriptome (Figure 1e) and each of the transcriptome datasets separately (short-read *de novo* assembly v1 – Figure 1a, short-read *de novo* assembly v2 – Figure 1b, and long-read Iso-Seq transcriptome – Figure 1d). b) Metrics determined for the *Pcitri*.v1 genome assembly, to which the transcriptome datasets were mapped.

**a) Transcriptome metrics**

| | Consolidated transcriptome | Long-read transcriptome | *de novo* v1 transcriptome | *de novo* v2 transcriptome |
|---|---|---|---|---|
| *Total number of assembled transcripts* | 440,881 | 72,445 | 317,306 | 134,464 |
| *Transcripts > 500* | 234,111 | 71,487 | 75,790 | 122,374 |
| *Transcripts >1000* | 168,571 | 63,478 | 44,334 | 81,127 |
| *Average length of assembled transcripts* | 1,356.41 | 2,358.59 | 633.64 | 2,180.98 |
| *Longest transcript* | 211,937 | 12,275 | 124,003 | 211,973 |
| *Total length* | 598,016,656 | 170,867,687 | 201,059,045 | 293,262,838 |
| *Transcript N50* | 2,908 | 2,800 | 1,515 | 3,495 |
| **Transcriptome alignment to *Pcitri*.v1 genome** | | | | |
| *Aligned* | 363,614 | 68,843 | 248,614 | 111,385 |
| *Uniquely aligned* | 315,563 | 55,010 | 229,305 | 91,910 |
| *Multipy aligned* | 6,828 | 35 | 6,962 | 651 |
| *Misassembly candidates reported by GMAP* | 41,058 | 13,759 | 12,297 | 19,038 |
| *Misassembly candidates reported by BLASTN* | 88,030 | 24,581 | 29,664 | 42,074 |
| *Misassemblies* | 22,983 | 8,801 | 5,270 | 11,052 |
| *Unaligned* | 77,267 | 3,602 | 68,692 | 23,079 |
| *Average aligned fraction* | 0.97 | 0.96 | 0.98 | 0.95 |
| *Average alignment length* | 1,196.85 | 2,120.38 | 596.42 | 1,952.25 |
| *Average blocks per alignment* | 5.93 | 12.44 | 2.95 | 8.64 |
| *Average block length* | 201.73 | 170.52 | 210.92 | 225.95 |
| *Average mismatches per transcript* | 8.31 | 6.17 | 6.80 | 9.71 |
| *NA50* | 2,546 | 2,550 | 1,310 | 3,002 |

| Assembly completeness (sensitivity) | | | | |
|---|---|---|---|---|
| *Database coverage* | 0.67 | 0.43 | 0.54 | 0.56 |
| *Duplication ratio* | 5.02 | 3.63 | 1.65 | 2.47 |
| *Average number of transcripts mapped to one isoform* | 5.20 | 3.64 | 2.52 | 2.68 |
| *50%-assembled genes* | 21,775 | 12,919 | 16,445 | 17,640 |
| *95%-assembled genes* | 16,687 | 10,806 | 10,887 | 13,392 |
| *50%-covered genes* | 22,664 | 12,988 | 18,039 | 19,015 |
| *95%-covered genes* | 17,929 | 11,082 | 12,718 | 14,218 |
| *50%-assembled isoforms* | 22,293 | 13,280 | 16,625 | 18,004 |
| *95%-assembled isoforms* | 16,976 | 11,005 | 10,949 | 13,558 |
| *50%-covered isoforms* | 23,203 | 13,352 | 18,244 | 18,404 |
| *95%-covered isoforms* | 18,258 | 11,298 | 12,789 | 14,403 |
| *50%-assembled exons* | 118,790 | 85,346 | 98,980 | 102,164 |
| *95%-assembled exons* | 113,418 | 83,603 | 91,952 | 98,760 |
| *Mean isoform assembly* | 0.81 | 0.96 | 0.69 | 0.84 |
| *Mean isoform coverage* | 0.84 | 0.93 | 0.74 | 0.86 |
| *Mean exon coverage* | 0.96 | 0.99 | 0.93 | 0.97 |
| *Average percentage of isoform 50%-covered exons* | 0.83 | 0.93 | 0.74 | 0.86 |
| *Average percentage of isoform 95%-covered exons* | 0.74 | 0.89 | 0.61 | 0.79 |
| **Assembly specificity** | | | | |
| *Unannotated* | 178,976 | 4,132 | 174,060 | 36,232 |
| *50%-matched* | 76,324 | 36,763 | 33,960 | 23,761 |
| *95%-matched* | 8,632 | 2,203 | 8,330 | 1,196 |
| *Mean fraction of transcript matched* | 0.23 | 0.59 | 0.14 | 0.27 |
| *Mean fraction of block matched* | 0.46 | 0.56 | 0.46 | 0.40 |
| *50%-matched blocks* | 0.46 | 0.56 | 0.47 | 0.40 |
| *95%-matched blocks* | 0.39 | 0.50 | 0.37 | 0.33 |
| *Matched length* | 157,253,815 | 72,465,708 | 41,811,301 | 64,516,908 |
| *Unmatched length* | 265,414,648 | 49,779,570 | 111,379,414 | 130,860,407 |

### b) Genome metrics

| ***Pcitri*.v1 *genome*** | |
|---|---|
| *Genes* | 40,620 |
| *Isoforms* | 42,260 |
| *Average length of all isoforms* | 1,104.78 |
| *Total length of all isoforms* | 46,687,951 |
| *Exons* | 180,975 |
| *Average exon length* | 257.98 |
| *Average number of exons per isoform* | 4.28 |
| *Maximal number of exons per isoform* | 117 |
| *Introns* | 138,715 |
| *Average intron length* | 853.25 |

**Table S2: Percent of short reads mapping to transcriptome sequences**. Illumina short-read samples used for *P. citri* short-read transcriptome assembly with SRA or GEO accession numbers and percentages of reads mapping to all three generated transcriptome datasets, counting uniquely and multimapped reads. The first seven samples were provided by Edinburgh University (E) and include samples from *P. citri* males (M) and females (F). Eight samples were sequenced in this study (S) and include samples from *P. citri* mated and virgin females (MF and VF, respectively). Reads from this study only were mapped back to v1 of short-read transcriptome, as it was assembled from those reads only.

| *Sample* | *Sample accession* | *Mapped to short-read v1* | *Mapped to short-read v2* | *Mapped to long-read* | *Average per sample* |
|---|---|---|---|---|---|
| EALL1 | / | / | 93.26% | 86.49% | **89.87%** |
| EF1 | SRR11260463 | / | 95.12% | 92.38% | **93.75%** |
| EF2 | SRR11260462 | / | 82.79% | 79.99% | **81.39%** |
| EF3 | SRR11260471 | / | 92.75% | 89.81% | **91.28%** |
| EM1 | SRR11260470 | / | 96.12% | 82.07% | **89.09%** |
| EM2 | SRR11260469 | / | 85.29% | 73.62% | **79.45%** |
| EM3 | SRR11260468 | / | 85.75% | 72.95% | **79.35%** |
| SMF08 | GSM5425814 | 74.95% | 78.53% | 62.37% | **71.95%** |
| SMF09 | GSM5425815 | 77.28% | 78.64% | 64.07% | **73.33%** |
| SMF10 | GSM5425816 | 77.88% | 81.10% | 70.48% | **76.49%** |
| SMF11 | GSM5425817 | 79.91% | 80.31% | 68.04% | **76.08%** |
| SVF02 | GSM5425810 | 78.15% | 80.84% | 68.72% | **75.90%** |
| SVF03 | GSM5425811 | 80.89% | 82.10% | 67.65% | **76.88%** |
| SVF05 | GSM5425812 | 84.30% | 86.23% | 75.74% | **82.09%** |
| SVF06 | GSM5425813 | 81.00% | 82.71% | 73.67% | **79.13%** |
| *Average per transcriptome set* | | **79.29%** | **85.44%** | **75.20%** | |

**Table S3: Differential expression of *P. citri* genes between virgin and mated females**, and

**Table S4: Selected IDS sequences**

are available as seperate files.

**Table S5: IDS activity measurements of *P. citri* candidates.** (a) Raw data for measurements of IDS enzymatic activity for candidate IDSs producing C10 or C 15 prenyl diphosphates, source data for Figure 3a, and (b) measurements of *trans*IDS5 mutants, source data for Figure 4a. Three measurements and their average are given for each protein and for each chain length. Values are given in mmol $s^{-1}$ $g^{-1}$.

a)

| | C10 | | | C15 | | | C10 average | C15 average |
|---|---|---|---|---|---|---|---|---|
| *trans*IDS5 | 133.71 | 137.93 | 139.84 | 169.16 | 201.27 | 209.93 | 137.16 | 193.46 |
| *trans*IDS3 | 22.74 | 24.00 | 21.96 | 2.57 | 2.58 | 2.40 | 22.90 | 2.51 |
| *trans*IDS11 | 0.66 | 0.60 | 0.55 | 0.27 | 0.25 | 0.23 | 0.60 | 0.25 |
| *trans*IDS2 | 0.60 | 0.60 | 0.56 | 0.32 | 0.29 | 0.24 | 0.59 | 0.28 |
| *trans*IDS17 | 0.09 | 0.09 | 0.10 | 0.04 | 0.03 | 0.04 | 0.09 | 0.04 |

b)

| | C10 | | | C15 | | | C10 average | C15 average |
|---|---|---|---|---|---|---|---|---|
| *trans*IDS5 -wt | 133.71 | 137.93 | 139.84 | 169.16 | 201.27 | 209.93 | 137.16 | 193.46 |
| D166N | 18.20 | 16.81 | 21.43 | 22.79 | 24.57 | 26.63 | 18.81 | 24.66 |
| D308N | 123.54 | 117.62 | 115.92 | 24.18 | 24.27 | 22.52 | 119.03 | 23.66 |
| D309N | 0.00 | 0.00 | 0.00 | 0.00 | 0.00 | 0.00 | 0.00 | 0.00 |
| D312N | 129.20 | 119.87 | 125.87 | 34.90 | 31.18 | 36.47 | 124.98 | 34.18 |
| K120A | 124.45 | 129.84 | 132.18 | 13.41 | 15.55 | 14.58 | 128.82 | 14.52 |
| K120E | 44.60 | 45.47 | 43.62 | 0.00 | 0.00 | 0.00 | 44.56 | 0.00 |
| K120Q | 120.59 | 118.66 | 120.46 | 11.94 | 11.95 | 12.04 | 119.90 | 11.98 |

**Table S6: Combined activity assays performed for detection of irregular IDS and terpene synthase activities.** All cloned IDS candidates were tested in pairwise combinations among each other (top part of the table) as well as in combination with other possible interactors: *P.citri* putative IDS regulatory subunits, g23689.tl, a putative *P. citri* pheromone binding protein (PBP); g36424.tl, *P.citri* isopentenyl diphosphate isomerase (IDI). For detection of terpene synthase activities, IDS-like candidates from *P. citri* were combined with IDS proteins reported to fulfil irregular coupling: chrysanthemyl diphosphate synthase from *Tanacetum cinerariifolium* (*Tc*CPPS) (Rivera et al. 2001), lavandulyl (*Li*LPPS) and cyclolavandulyl (*Ssp*CLPPS) diphosphate synthases from *Lavandula* × *intermedia* and *Streptomyces* sp. CL190, respectively (Demissie et al. 2013; Ozaki et al. 2014), and a mutant of neryl diphosphate synthase from *Solanum lycopersicum* (*Sl*NPPS-N88H) with increased irregular activity producing *trans*-planococcyl diphosphate (Gerasymenko et al. 2022).

| | *trans*IDS10 | *trans*IDS3 | *trans*IDS12 | *trans*IDS5 | *trans*IDS11 | *trans*IDS2 | *trans*IDS16 | *trans*IDS17 | *cis*IDS1 | *cis*IDS8 | |
|---|---|---|---|---|---|---|---|---|---|---|---|
| *trans*IDS10 | | X | X | X | X | X | X | X | X | X | Combinations between *P. citri* IDS candidates |
| *trans*IDS3 | | | X | X | X | X | X | X | X | X | |
| *trans*IDS12 | | | | X | X | X | X | X | X | X | |
| *trans*IDS5 | | | | | X | X | X | X | X | X | |
| *trans*IDS11 | | | | | | X | X | X | X | X | |
| *trans*IDS2 | | | | | | | X | X | X | X | |
| *trans*IDS16 | | | | | | | | X | X | X | |
| *trans*IDS17 | | | | | | | | | X | X | |
| *cis*IDS1 | | | | | | | | | | X | |
| *cis*IDS8 | | | | | | | | | | | |
| *trans*IDS4 | X | X | X | X | X | X | X | X | X | X | Combinations with regulatory subunits |
| *cis*IDS9 | X | X | X | X | X | X | X | X | X | X | |
| g23689.t1 (PBP) | X | X | X | X | X | X | X | X | X | X | Combinations with accessory proteins |
| g36424.t1 (IDI) | X | X | X | X | X | X | X | X | X | X | |
| *Tc*CPPS | X | X | X | X | X | X | X | X | X | X | Combinations with IDSs with irregular activity |
| *Li*LPPS | X | X | X | X | X | X | X | X | X | X | |
| *Ssp*CLPPS | X | X | X | X | X | X | X | X | X | X | |
| *Sl*NPPS-N88H | X | X | X | X | X | X | X | X | X | X | |

**Table S7: Input and output of the MEME motif search for *P. citri* sequences containing *trans*-IDS motifs.** Input sequences of FPPS-coding genes from different organisms used for MEME motif search (a) and output of MAST algorithm (b) with sequences from *Pcitri*.v1 genome, which contain the motifs detected by MEME. For the input sequences, organism names and Uniprot accession numbers are given, and for the target sequences, *Pcitri*.v1 gene model IDs and their E-values are given.

**a)**

| Organism | UniProt_acc |
|---|---|
| *Acromyrmex echinatior* | F4WXE7 |
| *Anthonomus grandis* | Q56CY8 |
| *Arabidopsis thaliana* | Q09152 |
| *Artemisia annua* | Q9ZPJ3 |
| *Artemisia spiciformis* | Q7XYS8 |
| *Bombyx mori* | Q95P28 |
| *Choristoneura fumiferana* | Q1XAB0 |
| *Culex quinquefasciatus* | B0VZA8 |
| *Dendroctonus jeffreyi* | Q56CY7 |
| *Drosophila melanogaster* | Q7KN61 |
| *Epicauta gorhami* | A0A075D844 |
| *Escherichia coli* | P22939 |
| *Gallus gallus* | P08836 |
| *Harpegnathos saltator* | E2B2E9 |
| *Ips pini* | Q58GE9 |
| *Mylabris cichorii* | A0A0U2D675 |
| *Myzus persicae* | B1PI49 |
| *Pediculus humanus corporis* | E0VQL3 |
| *Tanacetum cinerariifolium* | P0C565 |
| *Tetropium fuscum* | J9QQQ8 |
| *Tribolium castaneum* | D6WSE7 |

**b)**

| GeneID | Evalue |
|---|---|
| g32607.t1 | 5.30E-72 |
| g14484.t1 | 1.00E-63 |
| g21383.t1 | 8.10E-32 |
| g7366.t1 | 2.80E-25 |
| g34061.t1 | 1.50E-23 |
| g9107.t1 | 4.10E-18 |
| g2704.t1 | 4.20E-18 |
| g26044.t1 | 3.40E-10 |
| g21682.t1 | 8.30E-07 |
| g1822.t1 | 3.70E-05 |
| g14822.t1 | 1.30E-04 |
| g35512.t1 | 4.30E-01 |
| g35890.t1 | 7.30E+00 |
| g18384.t1 | 7.30E+00 |
| g4554.t1 | 7.90E+00 |
| g24024.t1 | 7.90E+00 |

**Table S8: Primers used for amplification of IDS sequences from *P. citri* cDNA and for introducing mutations into *trans*IDS5 sequence.** For cloning primers (a), the cloning sites for pMAL-c5X expression vector are underlined (NcoI in forward primer and BamHI in reverse primer), and for mutagenic primers (b), the mutated positions are underlined.

**a) Cloning primers**

| Primer pair | Target |
|---|---|
| Fw 5'-ACATCCATGGCGTGTTTGGGTTCG-3'<br>Rv 5'-ATGTGGATCCTACTTCAGACGGTTCAAC-3' | *trans*IDS2 |
| Fw 5'-ACATCCATGGCGAATTGTTCAAGTG-3'<br>Rv 5'-ATGATTCGGATCCTCAGAAATCCAC-3' | *trans*IDS4 |
| Fw 5'-GCACCATGGGTCCATTATTC-3'<br>Rv 5'-GATAGGATCCTTAATTACTCCGTTTG-3' | *trans*IDS5 |
| Fw 5'-ACATCCATGGATACTTTTGTTCGTTC-3'<br>Rv 5'-ATGTGGATCCTCATCTGCTATACTTTTTTATTAAATTC-3' | *trans*IDS11 |
| Fw 5'-ACATCCATGGTGAATTCGTTTCGAAG-3'<br>Rv 5'-ATCAGGATCCTTAAATTTCTCCCATCGC-3' | *trans*IDS12 |
| Fw 5'-ACATCCATGGATATGGAAAGTGG-3'<br>Rv 5'-TAGTGGATCCTCAAGTATTCTCTTCC-3' | *trans*IDS16 |
| Fw 5'-ACATCCATGGAAGACAAAGAAGC-3'<br>Rv 5'-ATGTGGATCCTTAATCGCTTTCGACC-3' | *trans*IDS17 |
| Fw 5'-TTGACCATGGCATCTGAACTACCAGCAC-3'<br>Rv 5'-TGAAGGATCCTCACTCCACGTGAATATATAATTTG-3' | *cis*IDS1 |
| Fw 5'-CTATCCATGGCGTCAAAAAATCAGCAACAAC-3'<br>Rv 5'-TCATGGATCCTTAGCTGTAATTCGGTGCTGTTC-3' | *cis*IDS8 |

### b) Mutagenic primers

| Primer pair | Mutation |
|---|---|
| Fw 5’- ATATTATGGATGGAGCTGAAACGAGAAG -3’<br>Rv 5’- CGTTCAATATTAAAAGAAATGCTTGAAGC -3’ | D166N |
| Fw 5’- ACGATTATTTGGACTGTTTTGG -3’<br>Rv 5’- TCTGTATTTGGAAATAATGTCCC-3’ | D308N |
| Fw 5’- TTGGACTGTTTTGGAGATGCAGATG -3’<br>Rv 5’- ATAATTGTCCTGTATTTGGAAATAATGTCC-3’ | D309N |
| Fw 5’- TTGGAGATGCAGATGAAATTGGTAAAATC -3’<br>Rv 5’- AACAGTTCAAATAATCGTCCTGTATTTGG-3’ | D312N |
| Fw 5’-AATCGTGGATTAGCTCTAGTCACCGC-3’<br>Rv 5’-TTTAGCTCCTCCGGGGACGTTATACTG-3’ | K120A |
| Fw 5’- AATCGTGGATTAGCTCTAGTCACCGC-3’<br>Rv 5’-TTTCTCTCCTCCGGGGACGTTATAC-3’ | K120E |
| Fw 5’- AATCGTGGATTAGCTCTAGTCACCGC-3’<br>Rv 5’- TTTCTGTCCTCCGGGGACGTTATAC -3’ | K120Q |

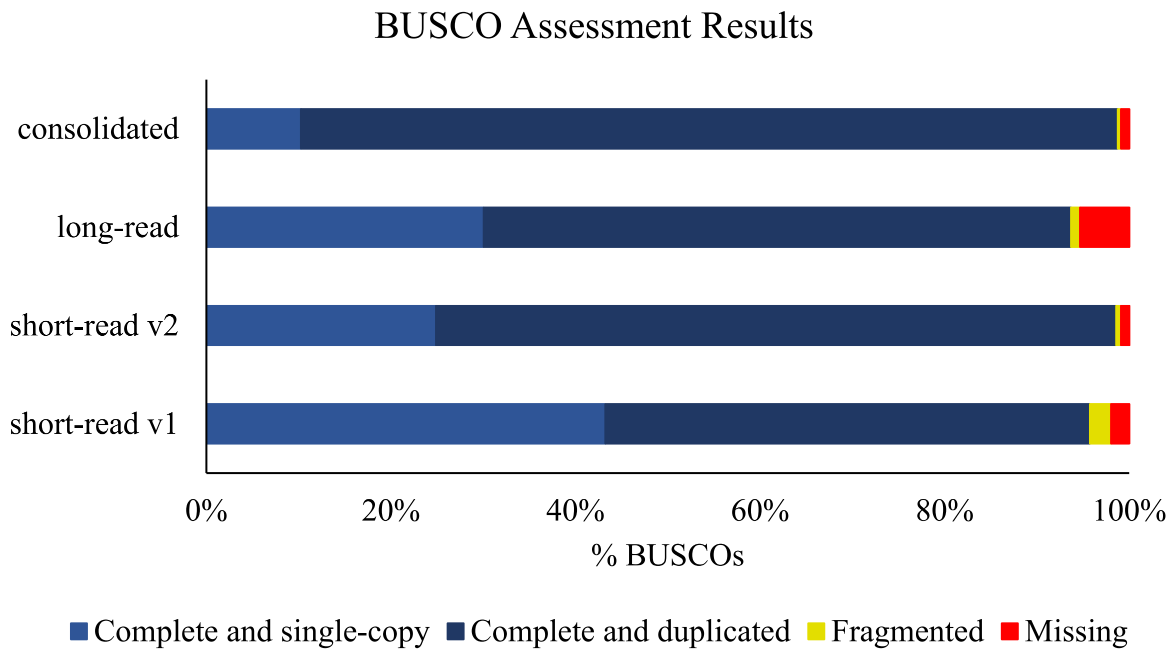

**Figure S1: BUSCO assessment of transcriptome completeness.** BUSCO was run for the consolidated transcriptome dataset (Figure 1e), long-read Iso-Seq dataset (Figure 1d) and both short-read *de novo* assembled datasets (Figure 1a and b). Percents of complete BUSCOs are shown in blue (light for single-copy and dark for duplicates) and for fragmented and missing BUSCOs in yellow and red, respectively. N=1367

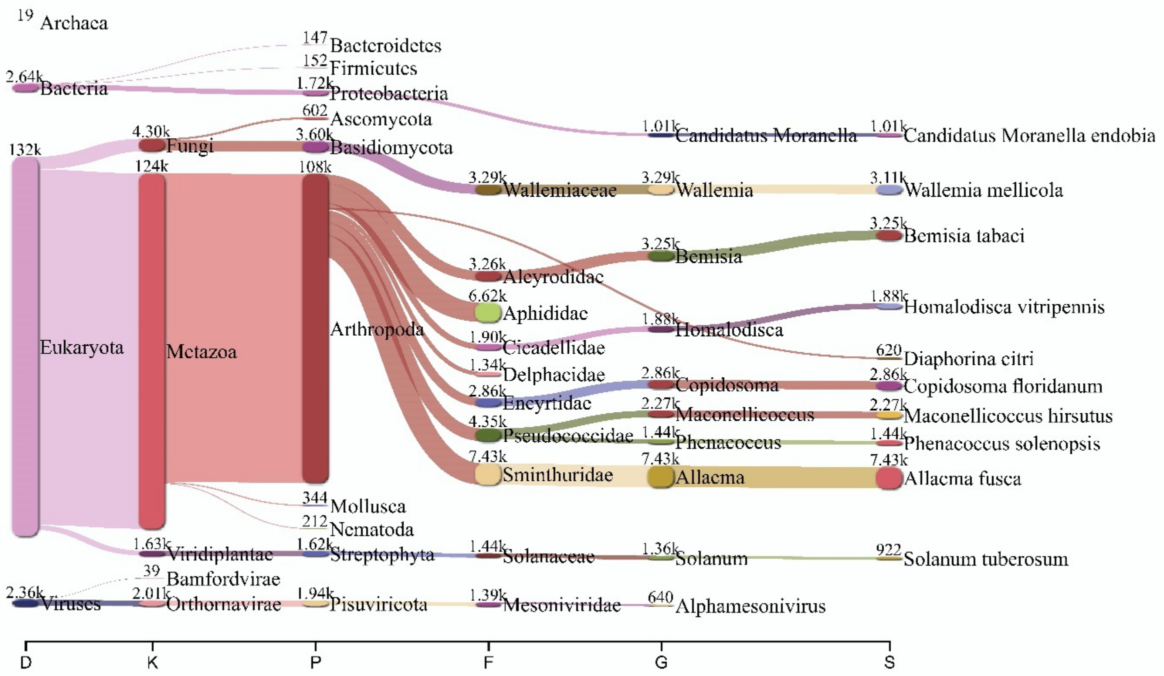

**Figure S2: Taxonomic classification of the *P. citri* transcriptome dataset.** Sankey plot visualisation of mmseqs2 classification of translated coding sequences from the consolidated *P. citri* transcriptome (Figure 1e). For each taxa, the number of assigned sequences is given. Markings on the x-axis: D - domain, K - kingdom, P - phylum, F - family, G - genus, S - species.

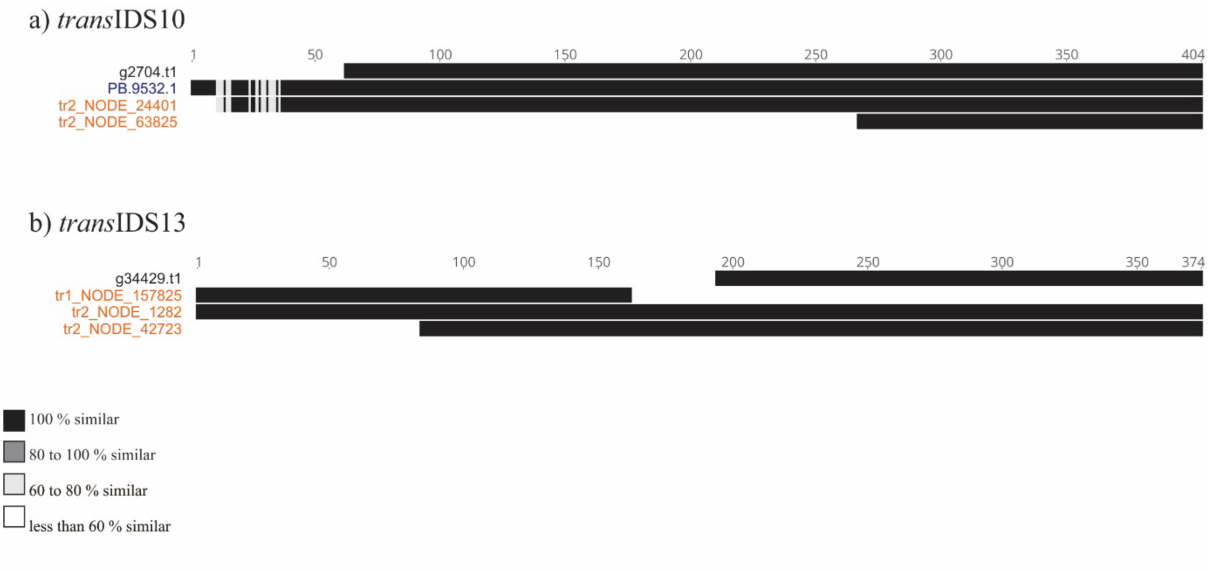

**Figure S3: Alignments of sequence resources for *trans*IDSl0 and *trans*IDS13.** Multiple sequence alignments were done in MEGAX and visualised with Geneious software. Color-coded similarity is given in the legend below. Names of sequences originating from the *Pcitri*.v1 genome, long-read, and short-read assemblies are given in black, blue and orange, respectively.

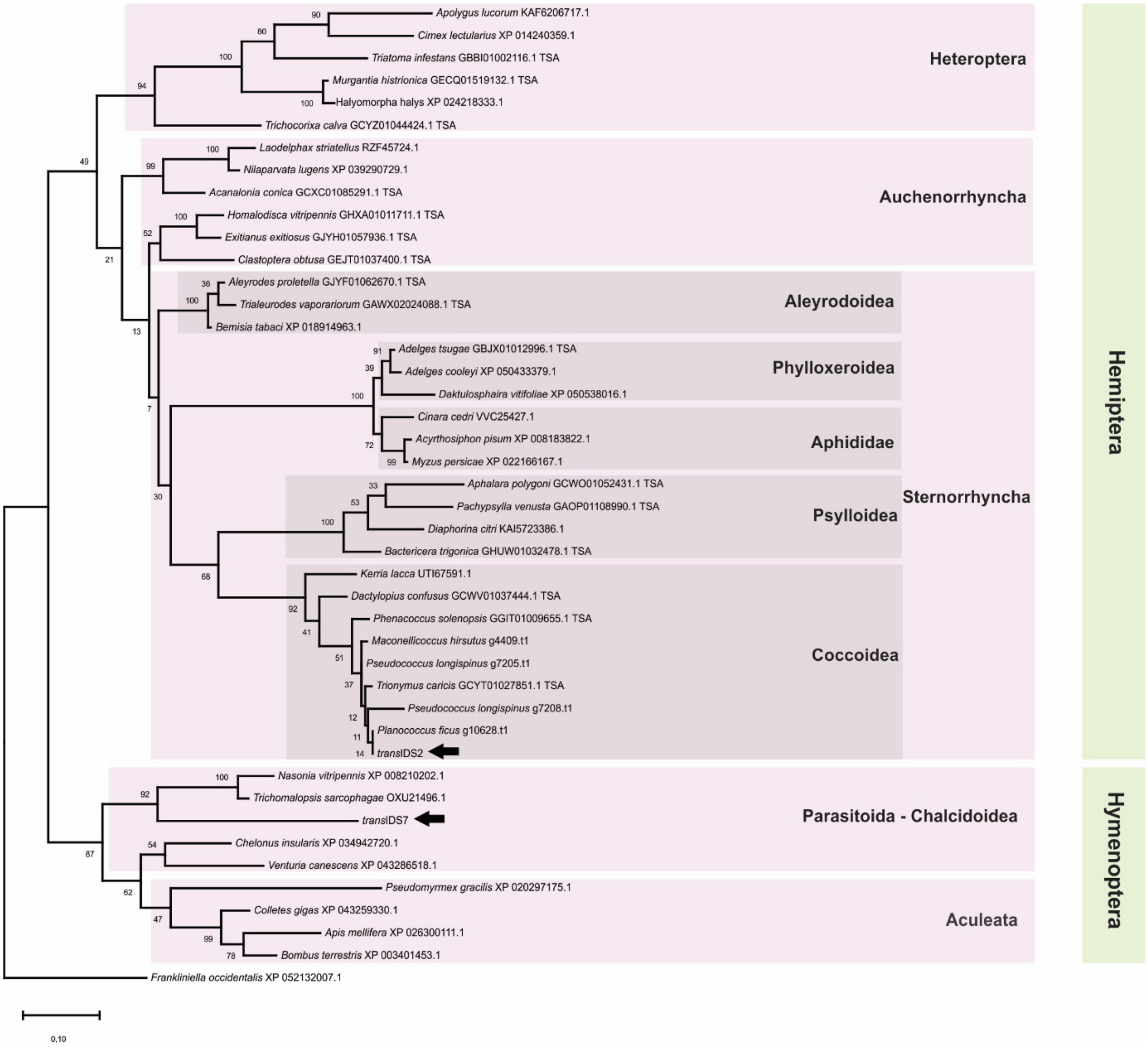

**Figure S4: Phylogenetic tree of putative DPPS subunit 1 sequences from selected species with *trans*IDS2 and *trans*IDS7.** The tree is drawn to scale, with branch lengths measured in the number of substitutions per site (scale on the bottom left) and bootstrap values given at nodes. This analysis involved 44 amino acid sequences with a total of 603 positions in the final dataset. Positions of candidate sequences from this study (*trans*IDS2 and *trans*IDS7) are marked with black arrows. For each sequence, species of origin and GenBank, TSA, or gene model ID are given. Taxonomic classification of included species is marked with coloured blocks.

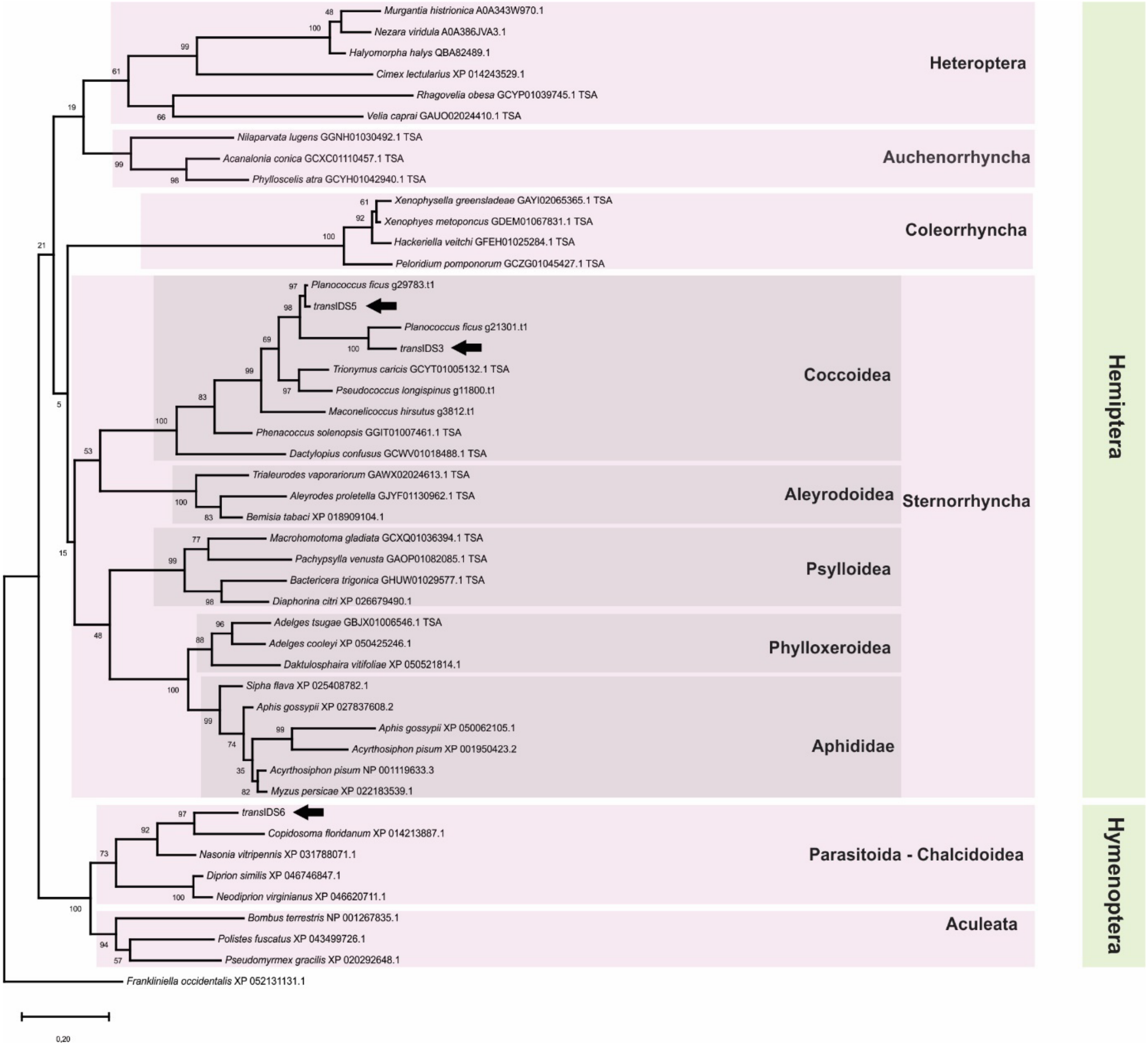

**Figure S5: Phylogenetic tree of putative FPPS sequences from selected species with *trans*IDS3, *trans*IDS5, and *trans*IDS6.** The tree is drawn to scale, with branch lengths measured in the number of substitutions per site (scale on the bottom left) and bootstrap values given at nodes. This analysis involved 47 amino acid sequences with a total of 525 positions in the final dataset. Positions of candidate sequences from this study (*trans*IDS3, *trans*IDS5, and *trans*IDS6) are marked with black arrows. For each sequence, species of origin and GenBank, TSA, or gene model ID are given. Taxonomic classification of included species is marked with coloured blocks.

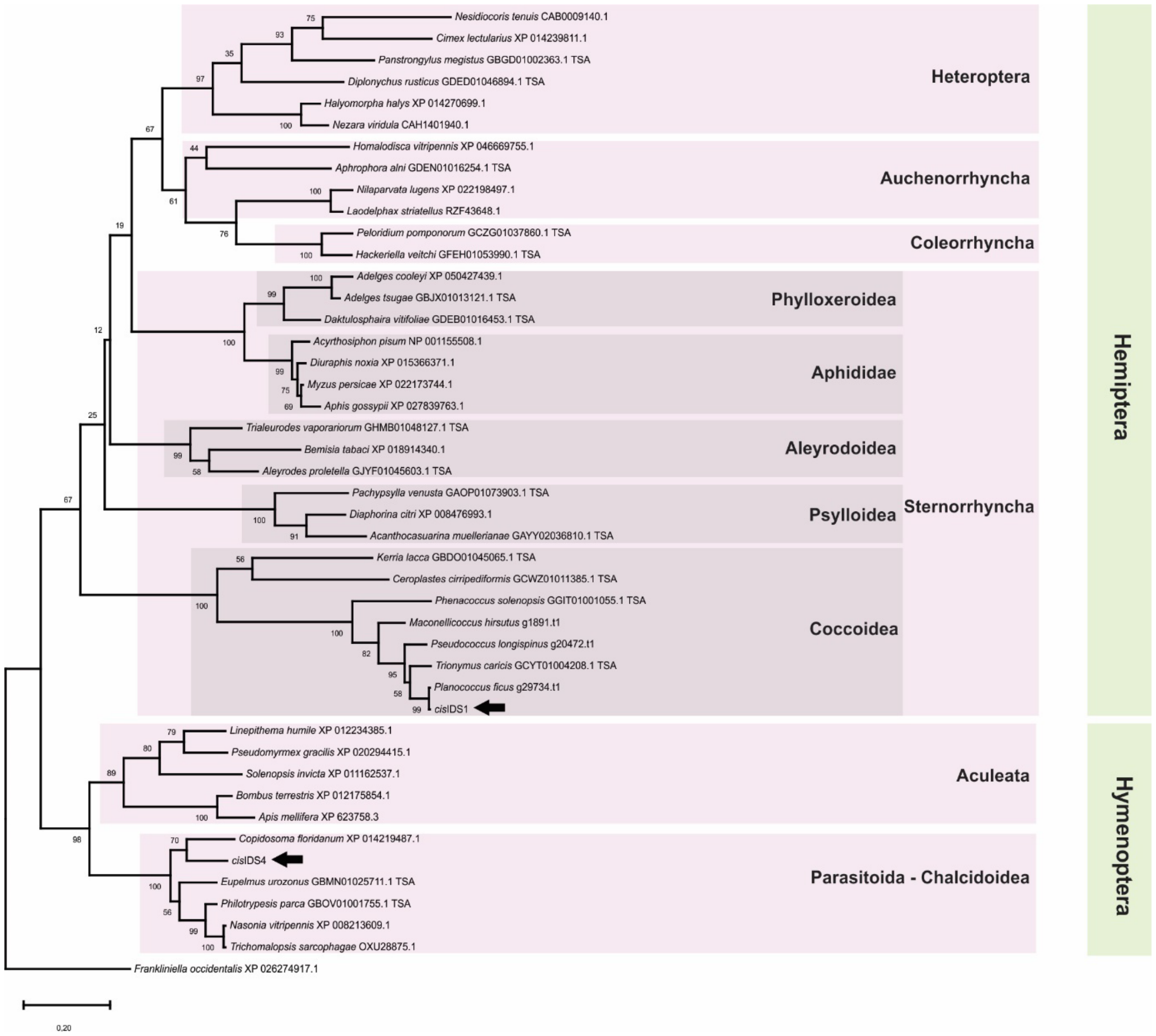

**Figure S6: Phylogenetic tree of putative DHPPS catalytic subunit sequences from selected species with *cis*IDS1, and *cis*IDS4.** The tree is drawn to scale, with branch lengths measured in the number of substitutions per site (scale on the bottom left) and bootstrap values given at nodes. This analysis involved 45 amino acid sequences with a total of 418 positions in the final dataset. Positions of candidate sequences from this study (*cis*IDS1, and *cis*IDS4) are marked with black arrows. For each sequence, species of origin and GenBank, TSA, or gene model ID are given. Taxonomic classification of included species is marked with coloured blocks.

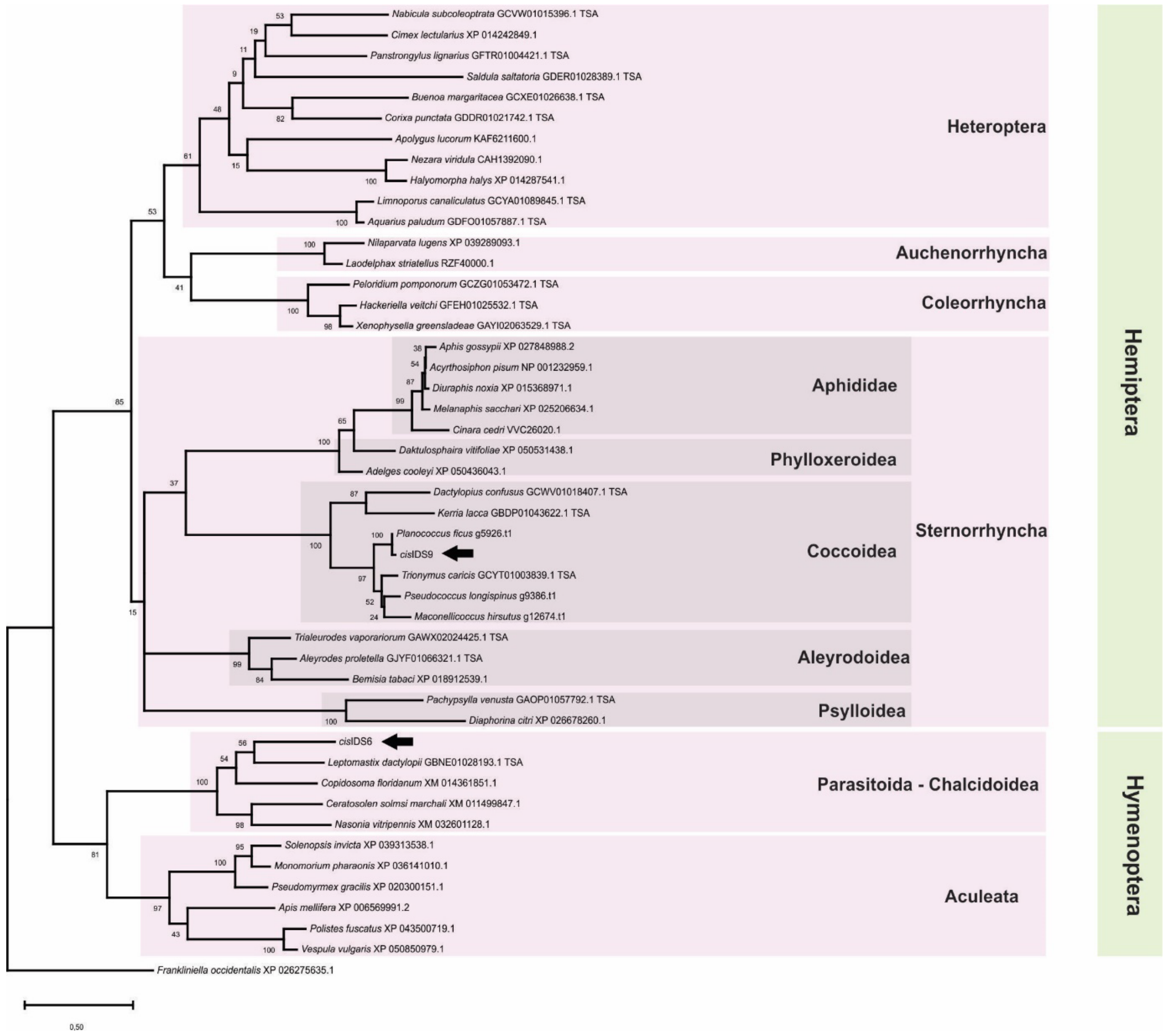

**Figure S7: Phylogenetic tree of putative DHPPS regulatory subunit sequences from selected species with *cis*IDS6, and *cis*IDS9.** The tree is drawn to scale, with branch lengths measured in the number of substitutions per site (scale on the bottom left) and bootstrap values given at nodes. The analysis involved 47 amino acid sequences with a total of 559 positions in the final dataset. Positions of candidate sequences from this study (*cis*IDS6, and *cis*IDS9) are marked with black arrows. For each sequence, species of origin and GenBank, TSA, or gene model ID are given. Taxonomic classification of included species is marked with coloured blocks.

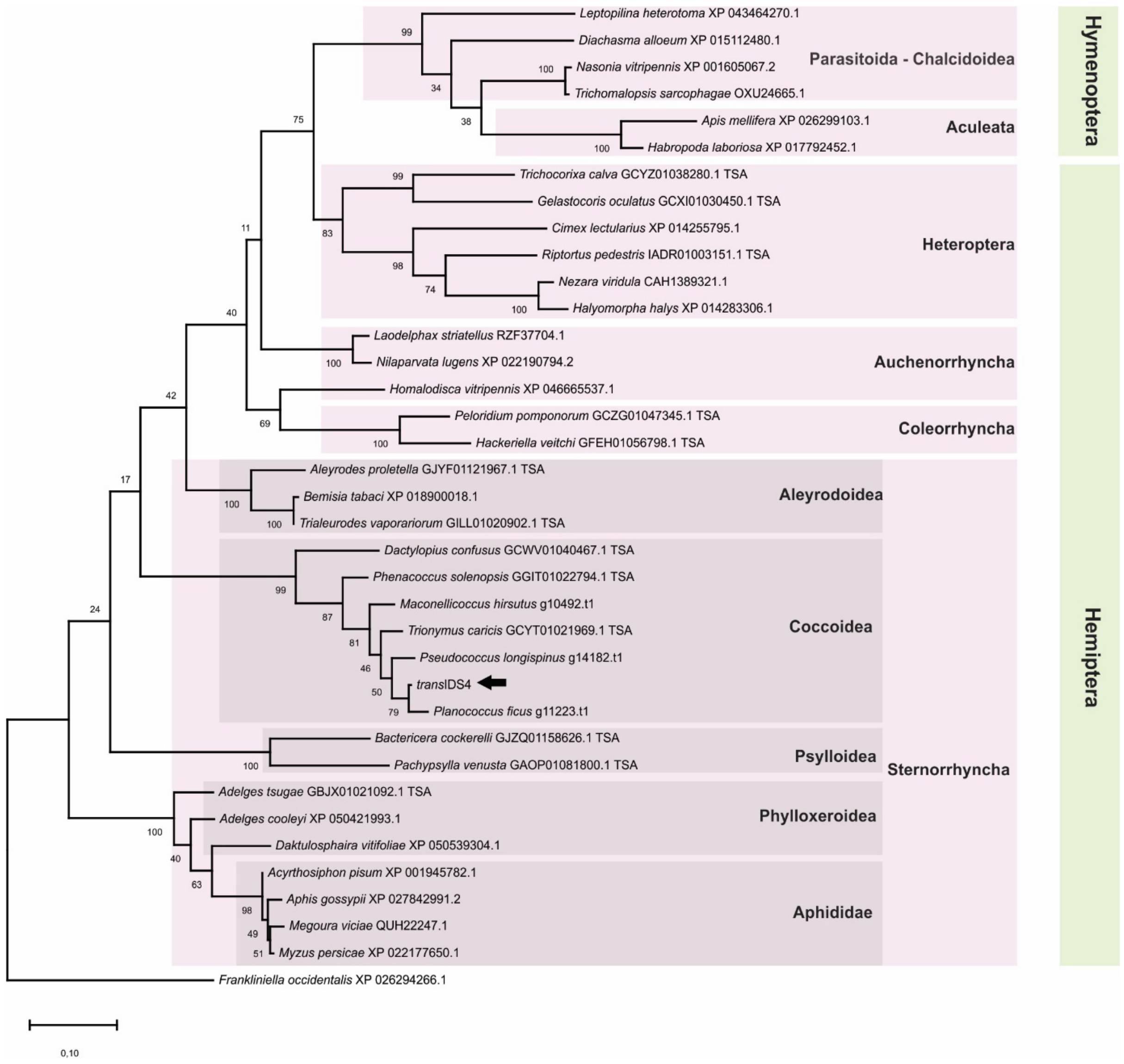

**Figure S8: Phylogenetic tree of putative DHPPS regulatory subunit sequences from selected species with *cis*IDS6, and *cis*IDS9.** The tree is drawn to scale, with branch lengths measured in the number of substitutions per site (scale on the bottom left) and bootstrap values given at nodes. The analysis involved 37 amino acid sequences with a total of 474 positions in the final dataset. Positions of candidate sequences from this study (*cis*IDS6, and *cis*IDS9) are marked with black arrows. For each sequence, species of origin and GenBank, TSA, or gene model ID are given. Taxonomic classification of included species is marked with coloured blocks.

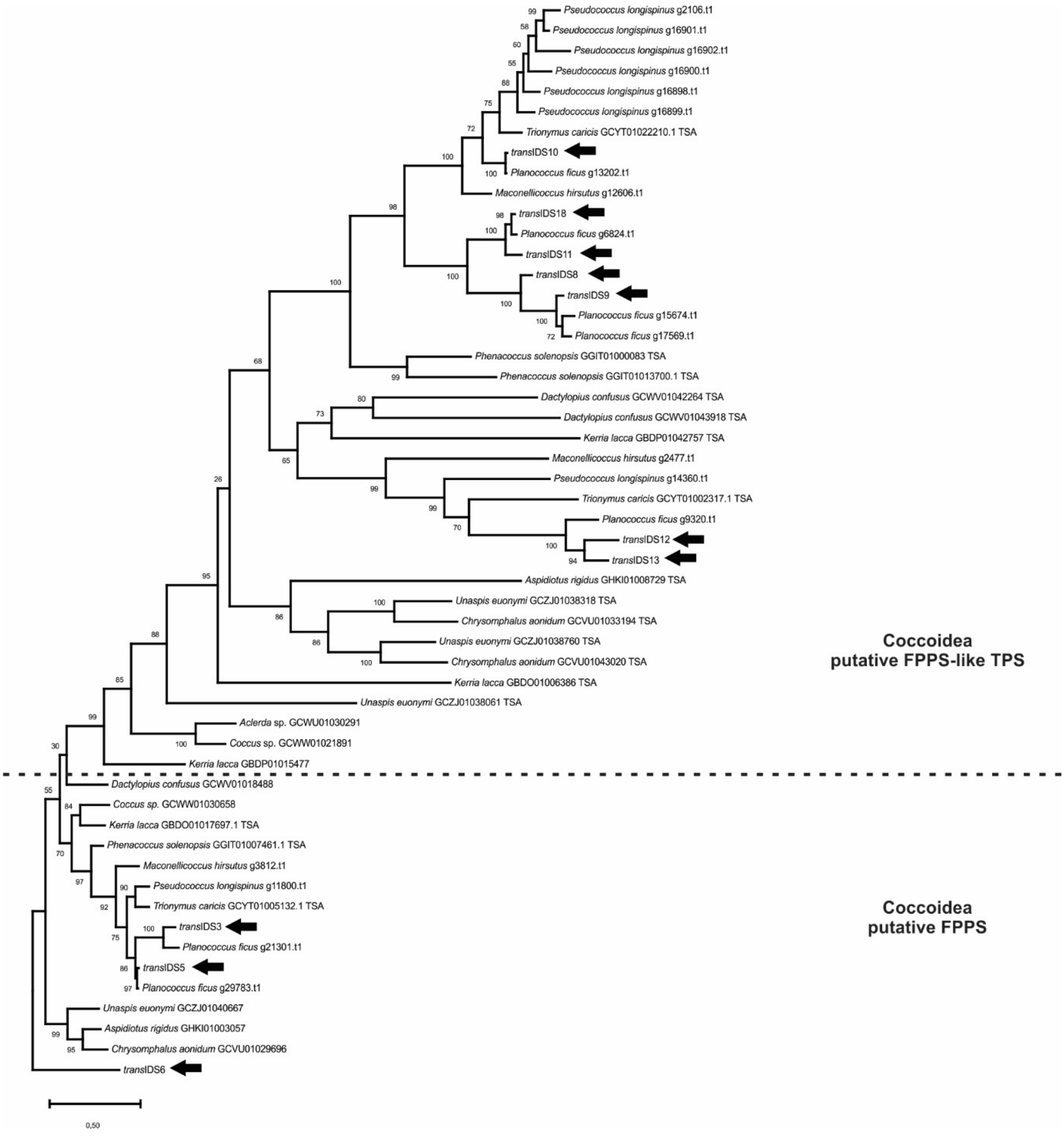

**Figure S9: Phylogenetic tree of putative FPPS and FPPS-like sequences from Coccoidea species.** The tree is drawn to scale, with branch lengths measured in the number of substitutions per site (scale on the bottom left) and bootstrap values given at nodes. The analysis involved 53 amino acid sequences with a total of 826 positions in the final dataset. Positions of candidate sequences from this study (*trans*IDS3, *trans*IDS5, *trans*IDS6, *trans*IDS8, *trans*IDS9, *trans*IDS10, *trans*IDS11, *trans*IDS12, *trans*IDS13, *trans*IDS18) are marked with black arrows. For each sequence, species of origin and GenBank, TSA, or gene model ID are given. Taxonomic classification of included species is marked with coloured blocks. The delineation between the FPPS and FPPS-like sequences is based on Rebholz et al., 2023.

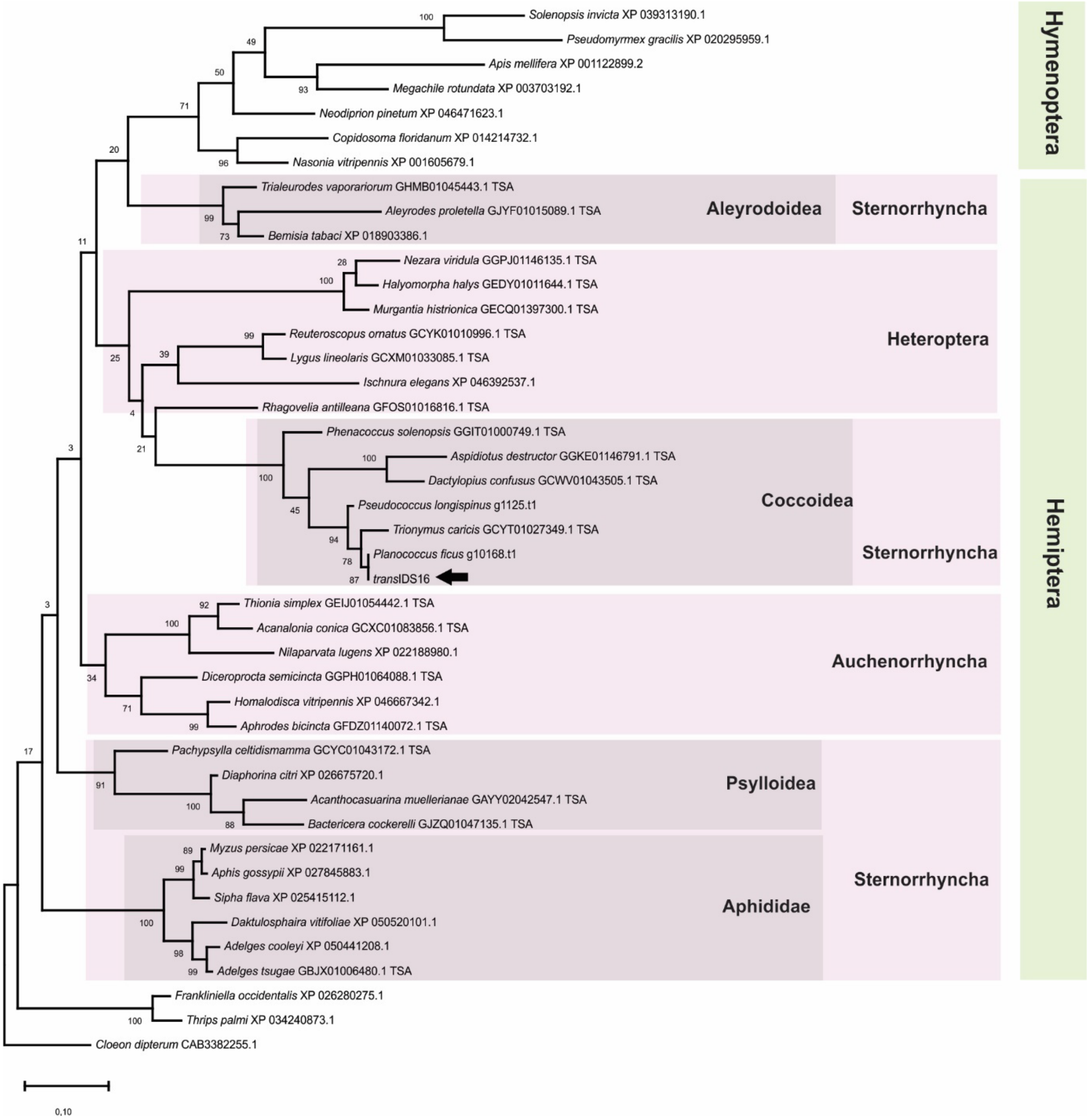

**Figure S10: Phylogenetic tree of putative GGPPS sequences from selected species with *trans*IDS16.** The tree is drawn to scale, with branch lengths measured in the number of substitutions per site (scale on the bottom left) and bootstrap values given at nodes. The analysis involved 43 amino acid sequences with a total of 400 positions in the final dataset. Positions of candidate sequences from this study (*trans*IDS16) are marked with black arrows. For each sequence, species of origin and GenBank, TSA, or gene model ID are given. Taxonomic classification of included species is marked with coloured blocks.

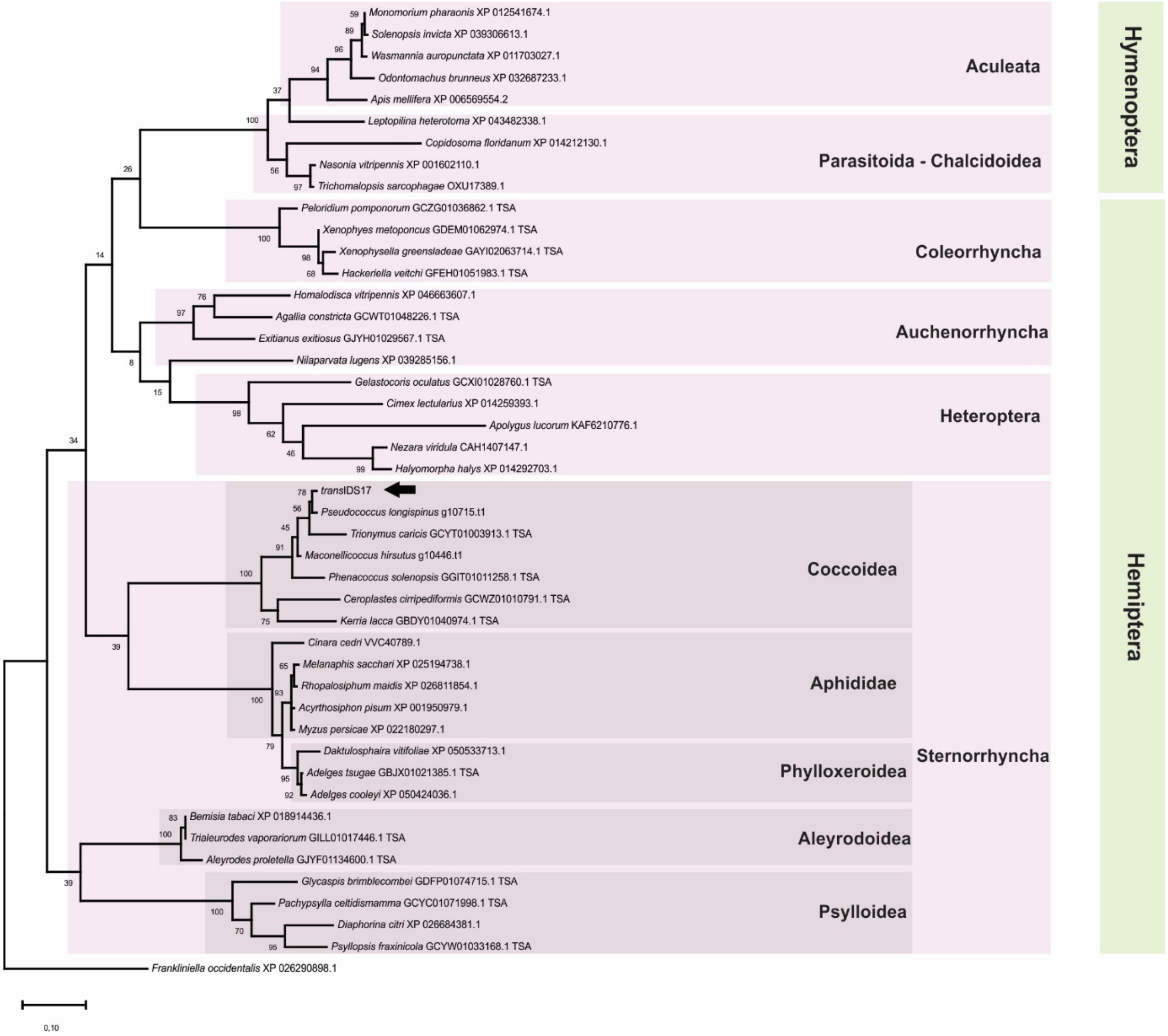

**Figure S11: Phylogenetic tree of putative otubain-like sequences from selected species with *trans*IDS17.** The tree is drawn to scale, with branch lengths measured in the number of substitutions per site (scale on the bottom left) and bootstrap values given at nodes. The analysis involved 45 amino acid sequences with a total of 344 positions in the final dataset. Positions of candidate sequences from this study (*trans*IDS17) are marked with black arrows. For each sequence, species of origin and GenBank, TSA, or gene model ID are given. Taxonomic classification of included species is marked with coloured blocks.

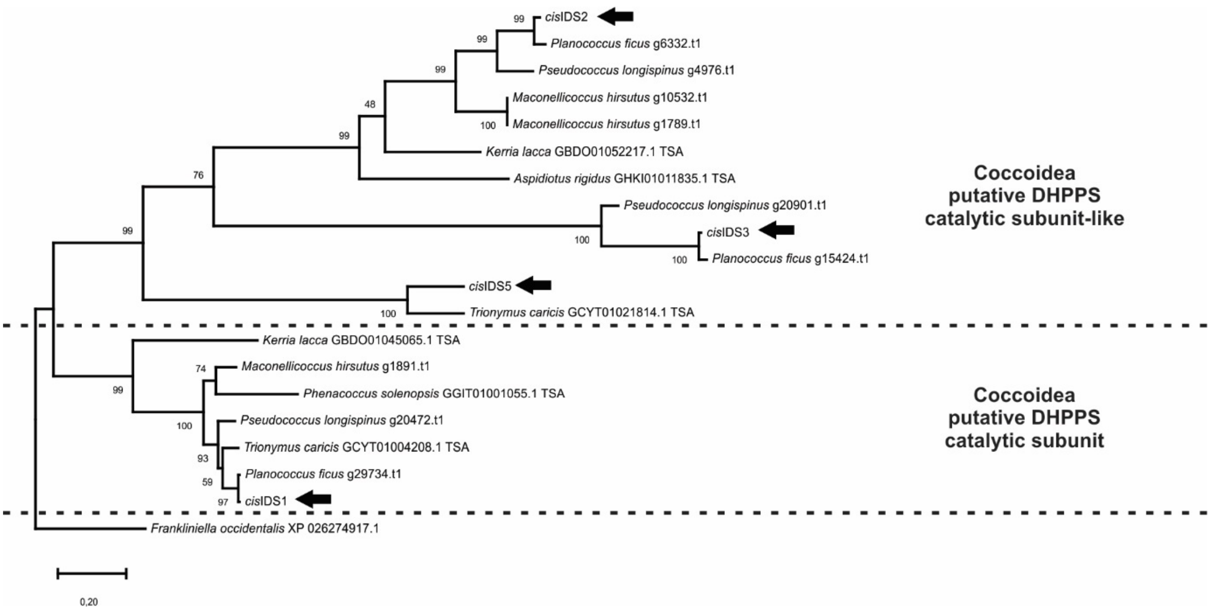

**Figure S12: Phylogenetic tree of putative DHPPS and DHPPS-like sequences from selected Coccoidea species.** The tree is drawn to scale, with branch lengths measured in the number of substitutions per site (scale on the bottom left) and bootstrap values given at nodes. The analysis involved 45 amino acid sequences with a total of 344 positions in the final dataset. Positions of candidate sequences from this study (*cis*IDS1, *cis*IDS2, *cis*IDS3, *cis*IDS5) are marked with black arrows. For each sequence, species of origin and GenBank, TSA, or gene model ID are given. Taxonomic classification of included species is marked with coloured blocks.

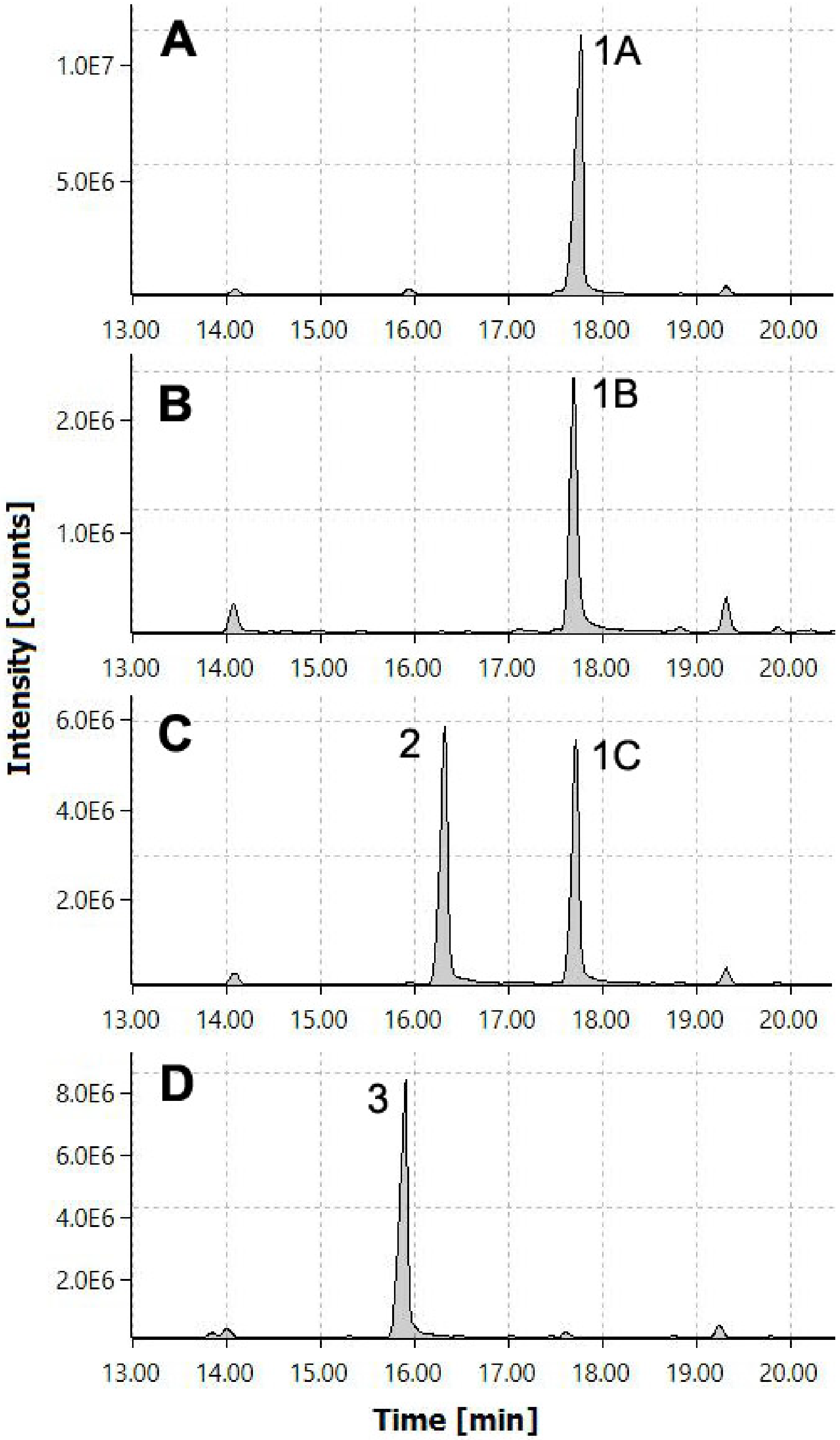

**Figure S13: Identification of regular monoterpenes synthesised by *trans*-IDS enzymes from *P. citri*.** The proteins were incubated with IPP and DMAPP, and dephosphorylated products were analysed by GC-MS. (A) Standard GPP (dephosphorylated) – geraniol (peak 1A); (B) *trans*IDS5 – geraniol (peak 1B); (C) *trans*IDS3 – geraniol (peak 1C) and iso-geraniol (peak 2); (D) standard NPP (dephosphorylated) – nerol (peak 3). EI-MS data of the peaks are shown in Figures S14 and S15.

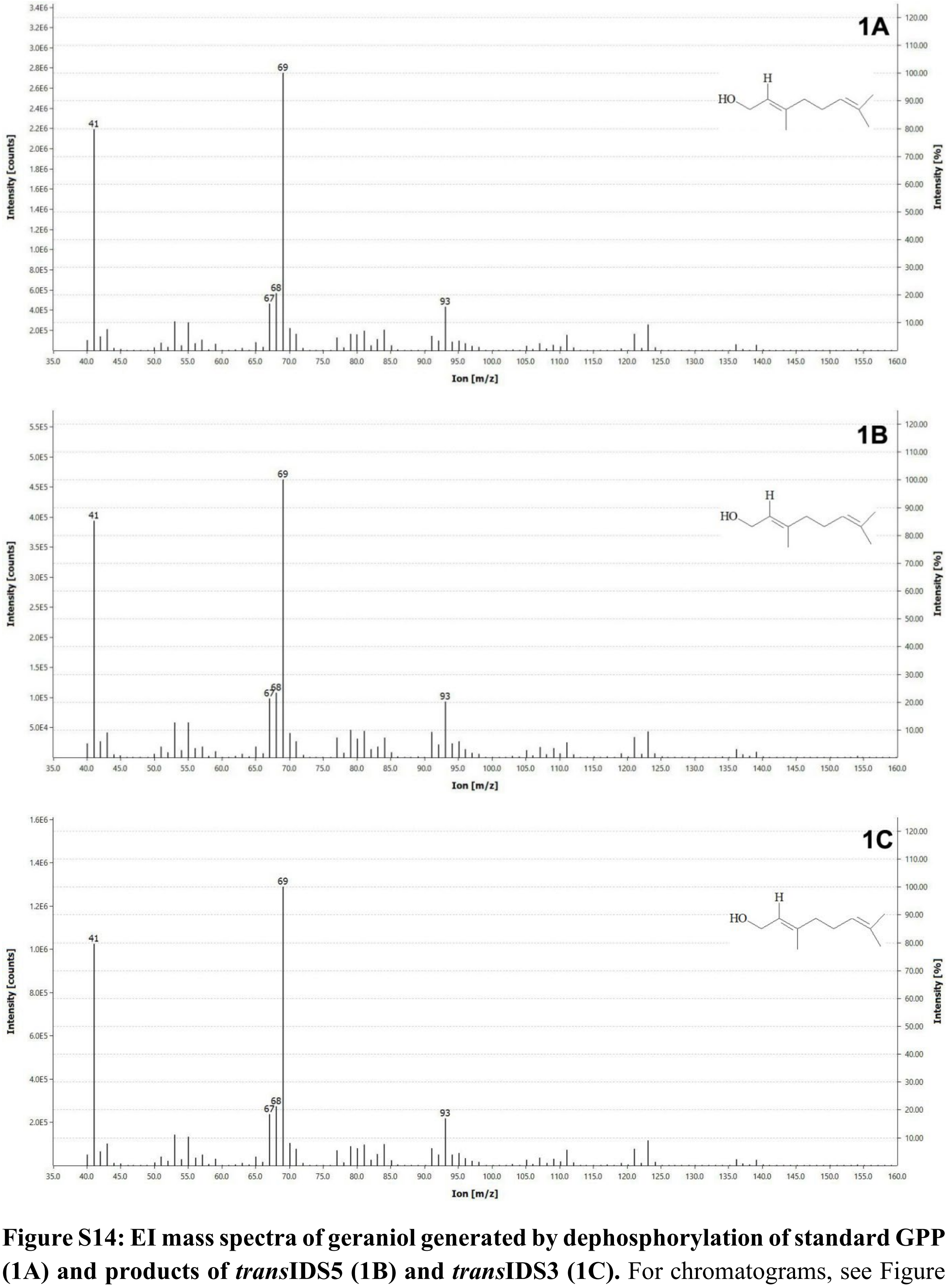

**Figure S14: EI mass spectra of geraniol generated by dephosphorylation of standard GPP (1A) and products of *trans*IDS5 (1B) and *trans*IDS3 (1C).** For chromatograms, see Figure S13.

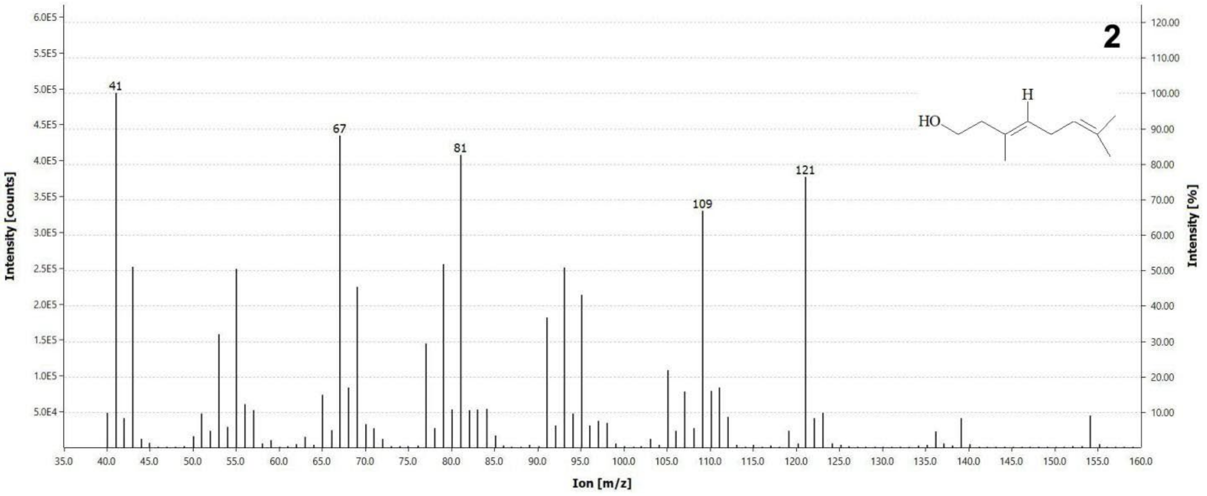

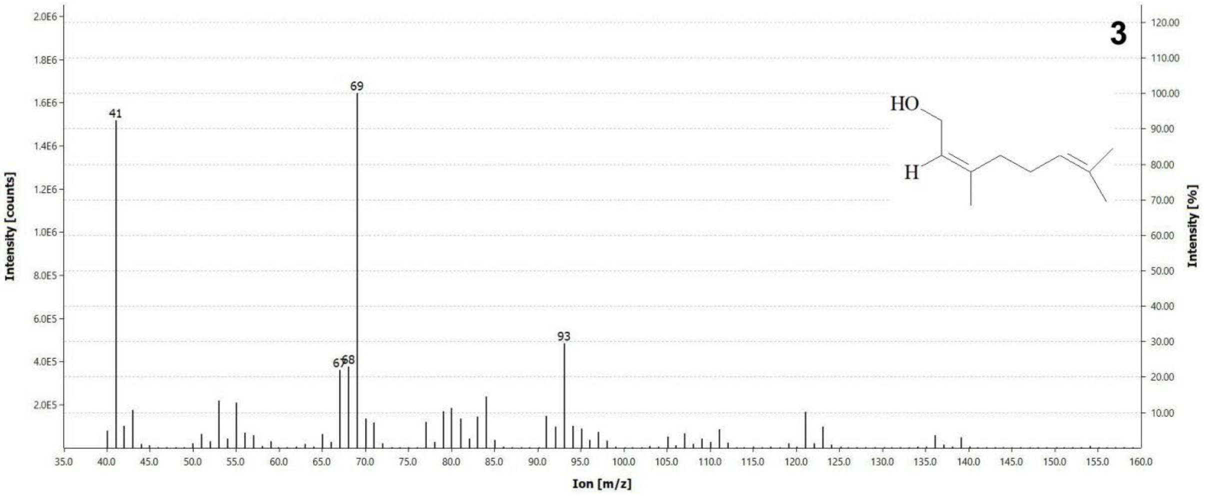

**Figure S15: EI mass spectra of dephosphorylated derivatives of product of *trans*IDS3 (2, iso-geraniol) and standard NPP (3, nerol).** For chromatograms, see Figure S13.

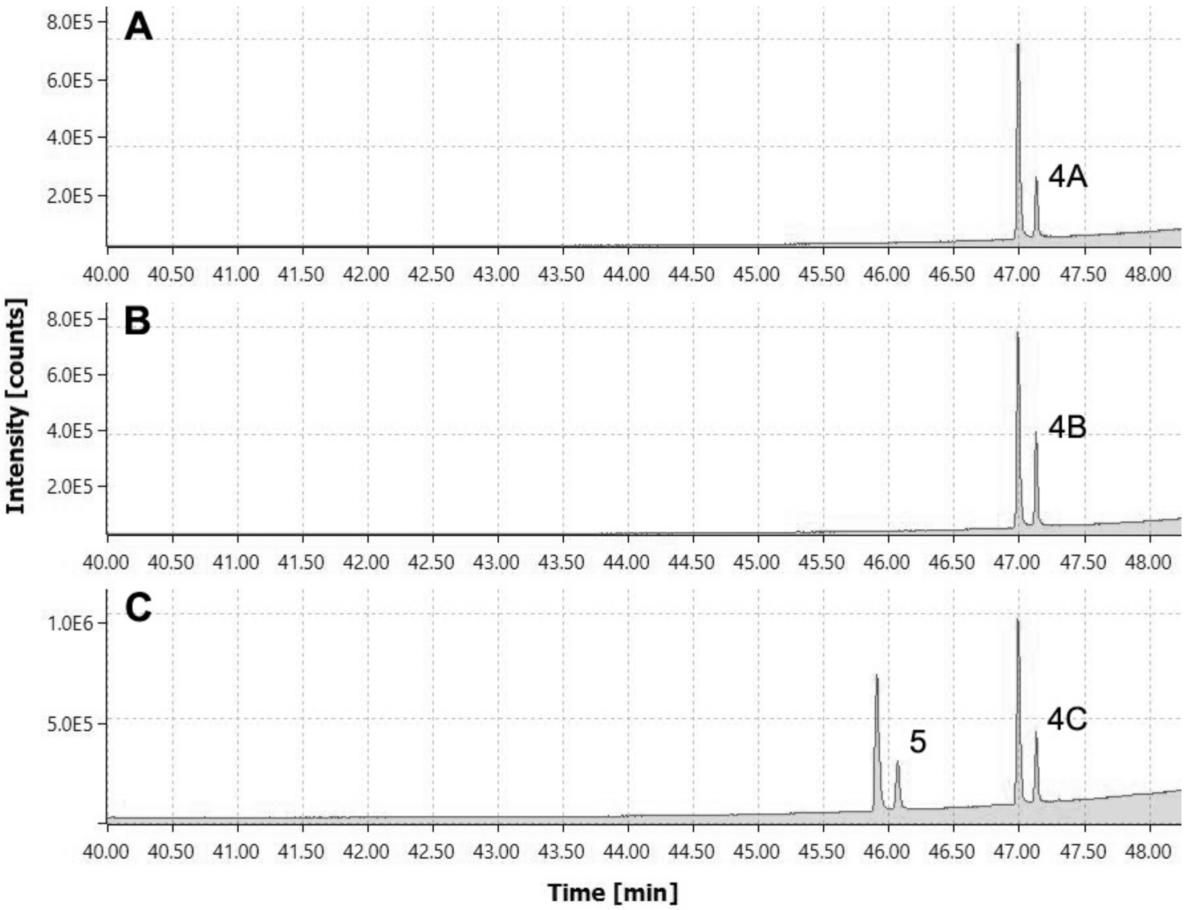

**Figure S16: Identification of regular sesquiterpenes synthesised by *trans*-IDS enzymes from *P. citri*.** The proteins were incubated with IPP and DMAPP, and dephosphorylated products were analysed by GC-MS. (A) FPPS from *Tanacetum cinerariifolium* (known to produce *trans*-farnesyl diphosphate) – *trans*-farnesol (peak 4A); (B) *trans*IDS5 – *trans*-farnesol (peak 4B); (C) *trans*IDS3 – *trans*-farnesol (peak 4C) and farnesol isomer (peak 5). EI-MS data of the peaks are shown in Figures S17 and S18.

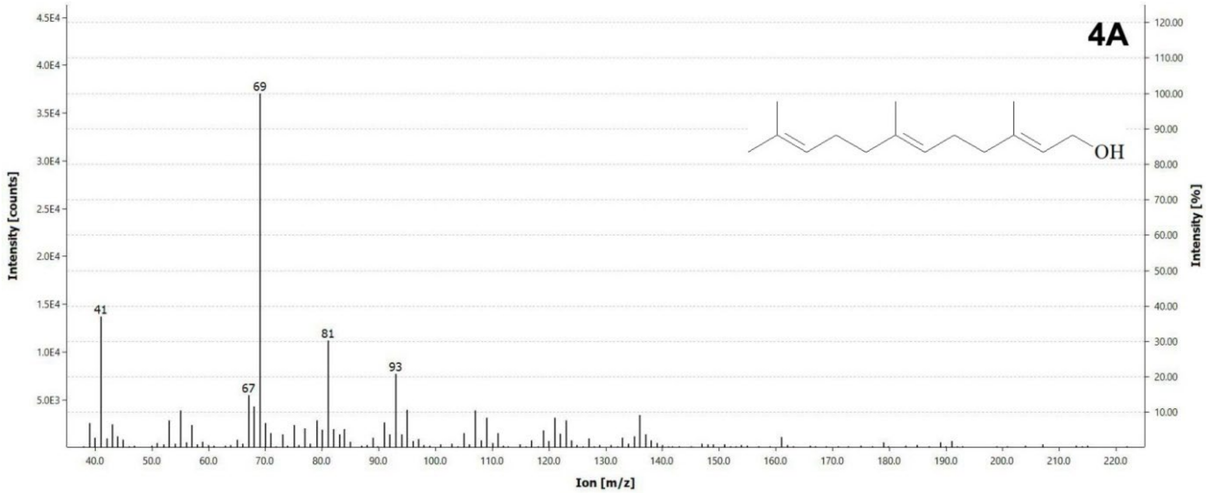

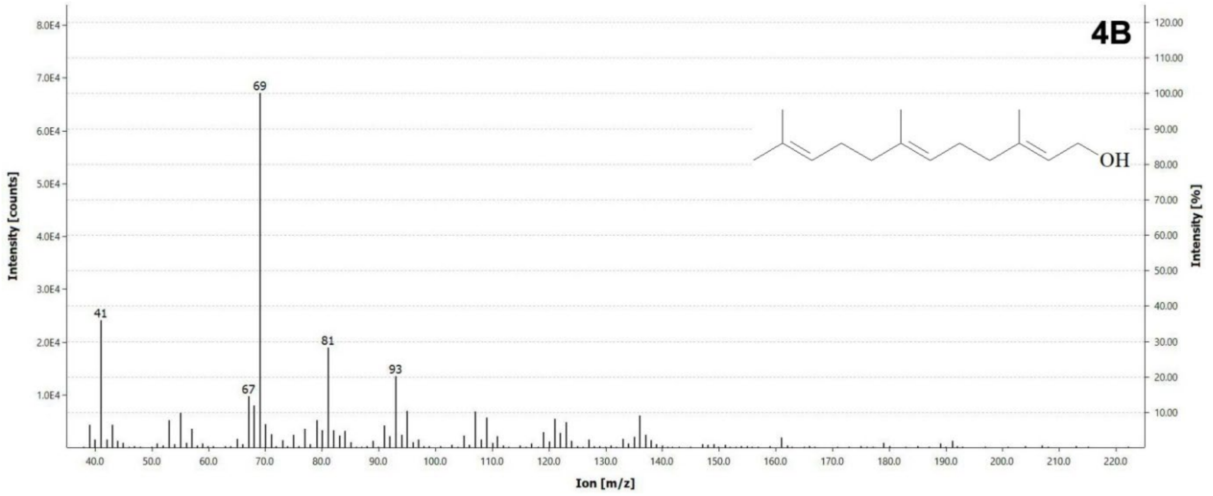

**Figure S17: EI mass spectra of *trans*-farnesol generated by dephosphorylation of products of FPPS from *Tanacetum cinerariifolium* known to produce *trans*-farnesyl diphosphate (4A) and *trans*IDS5 (4B).** For chromatograms, see Figure S16.

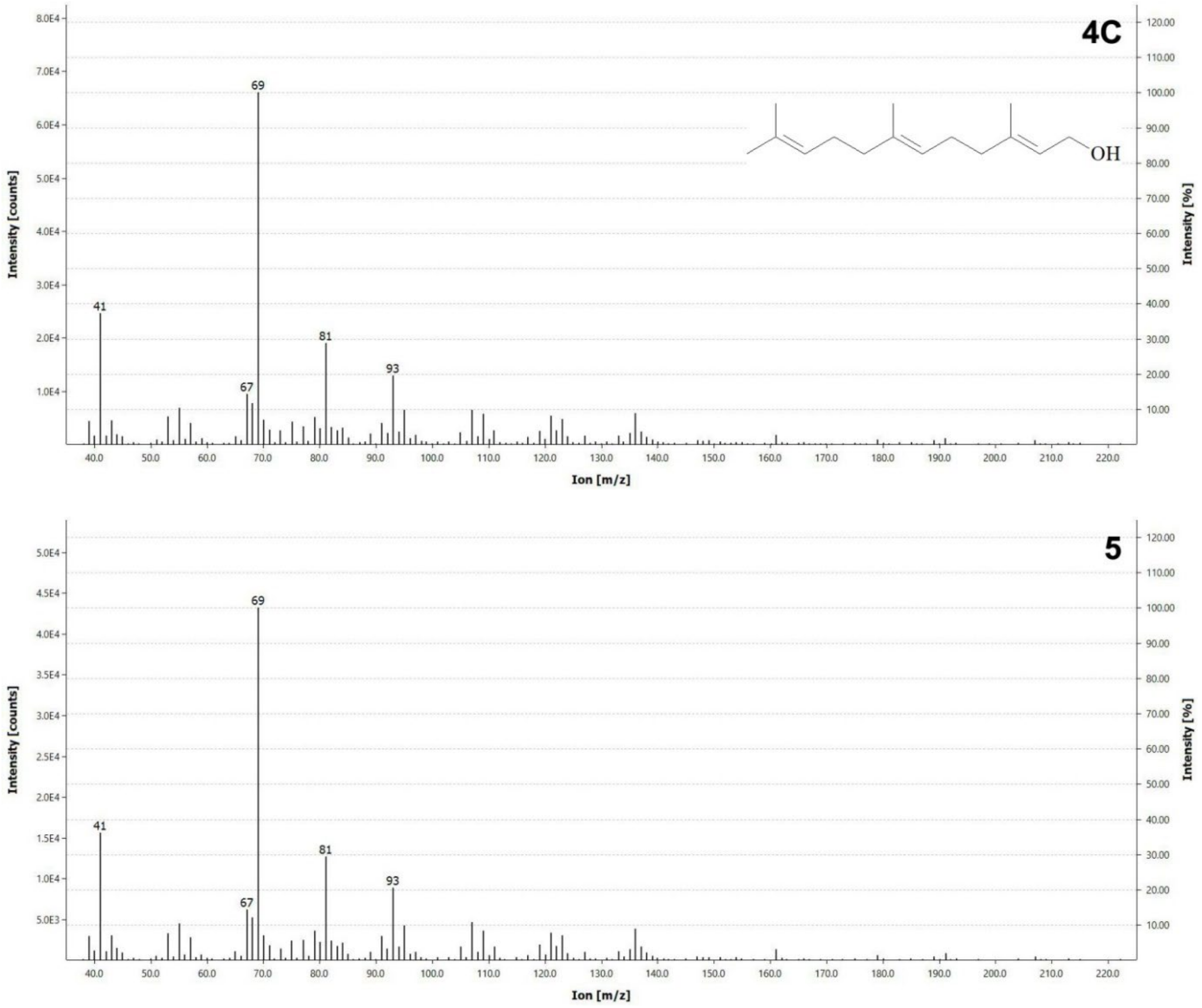

**Figure S18: EI mass spectra of dephosphorylated products of *trans*IDS3, *trans*-farnesol (4C) and farnesol isomer (5).** For chromatograms, see Figure S16.

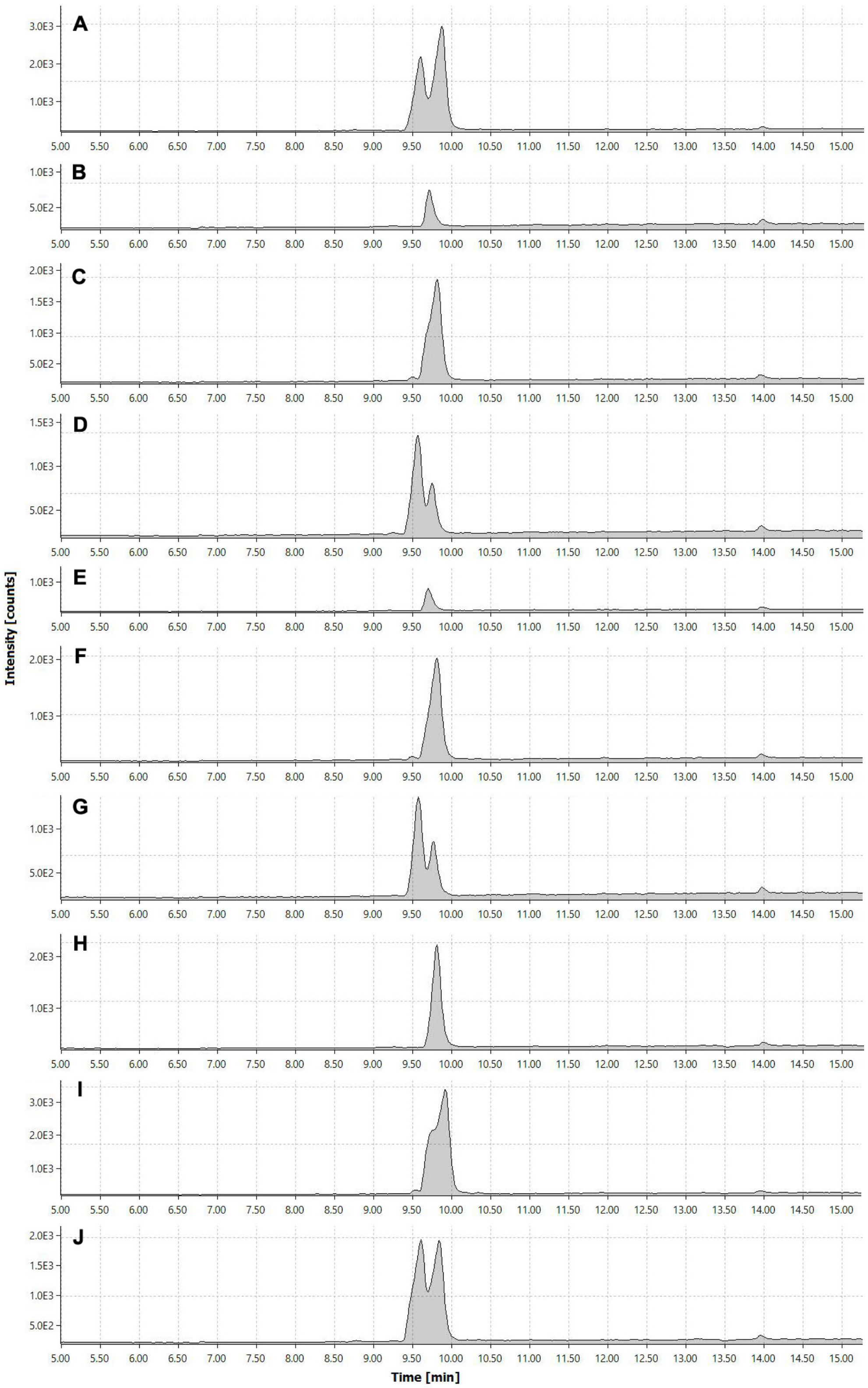

**Figure S19: Identification of regular monoterpene diphosphates synthesised by *trans*-IDS enzymes from *P. citri*.** The proteins were incubated with IPP and DMAPP, and the products were analysed by LC-MS. (A) Standard NPP and GPP; (B) product of *trans*IDS2; (C) product of *trans*IDS2 and GPP; (D) product of *trans*IDS2 and NPP; (E) product of *trans*IDS11; (F) product of *trans*IDS11 and GPP; (G) product of *trans*IDS11 and NPP; (H) product of *trans*IDS17; (I) product of *trans*IDS17 and GPP; (J) product of *trans*IDS17 and NPP.

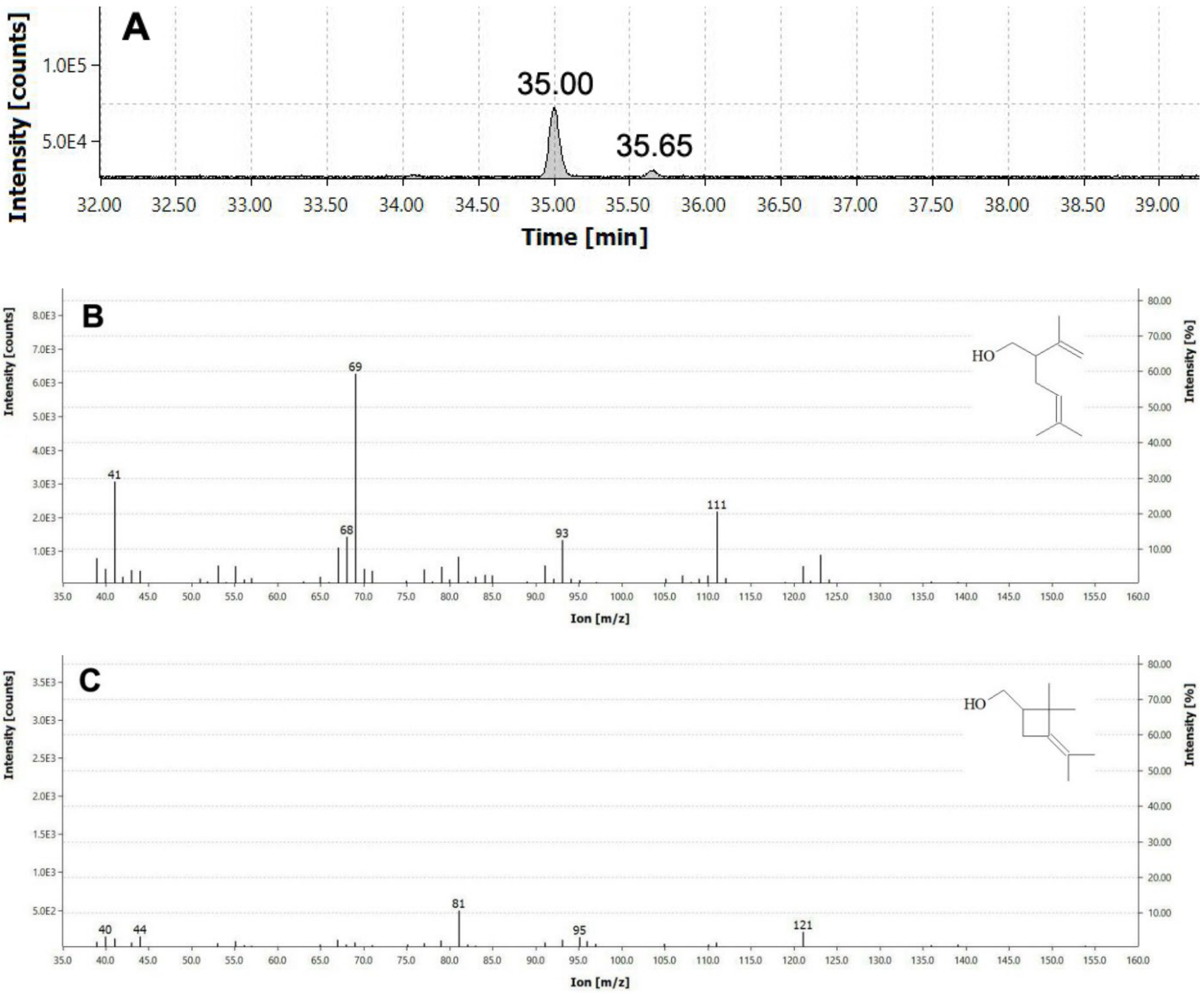

**Figure S20. Identification of irregular monoterpenes synthesised by *trans*IDS5.** The protein was incubated with DMAPP only, and dephosphorylated products were analysed by GC-MS. (A) TIC chromatogram with two product peaks; (B) EI mass spectrum of peak at RT 35.00 min (lavandulol); (C) EI mass spectrum of peak at RT 35.65 min (maconelliol).
